# Proportional intracranial volume correction differentially biases behavioral predictions across neuroanatomical features and populations

**DOI:** 10.1101/2022.03.15.483970

**Authors:** Elvisha Dhamala, Leon Qi Rong Ooi, Jianzhong Chen, Ru Kong, Kevin M. Anderson, Rowena Chin, B.T. Thomas Yeo, Avram J. Holmes

## Abstract

Individual differences in brain anatomy can be used to predict variability in cognitive function. Most studies to date have focused on broad population-level trends, but the extent to which the observed predictive features are shared across sexes and age groups remains to be established. While it is standard practice to account for intracranial volume (ICV) using proportion correction in both regional and whole-brain morphometric analyses, in the context of brain-behavior predictions the possible differential impact of ICV correction on anatomical features and subgroups within the population has yet to be systematically investigated. In this work, we evaluate the effect of proportional ICV correction on sex-independent and sex-specific predictive models of individual cognitive abilities across multiple anatomical properties (surface area, gray matter volume, and cortical thickness) in healthy young adults (Human Connectome Project; n=1013, 548 females) and typically developing children (Adolescent Brain Cognitive Development study; n=1823, 979 females). We demonstrate that ICV correction generally reduces predictive accuracies derived from surface area and gray matter volume, while increasing predictive accuracies based on cortical thickness in both adults and children. Furthermore, the extent to which predictive models generalize across sexes and age groups depends on ICV correction: models based on surface area and gray matter volume are more generalizable without ICV correction, while models based on cortical thickness are more generalizable with ICV correction. Finally, the observed neuroanatomical features predictive of cognitive abilities are unique across age groups regardless of ICV correction, but whether they are shared or unique across sexes (within age groups) depends on ICV correction. These findings highlight the importance of considering individual differences in ICV, and show that proportional ICV correction does not remove the effects of cranium volumes from anatomical measurements and can introduce ICV bias where previously there was none. ICV correction choices affect not just the strength of the relationships captured, but also the conclusions drawn regarding the neuroanatomical features that underlie those relationships.

## Introduction

A primary goal of research in the brain sciences is to establish the relationship between neurobiological features and behavioral traits, allowing for both the understanding and prediction of individual differences across health and disease (Yarkoni and Westfall 2017; Kohoutová et al. 2020; Bzdok and Yeo 2017). While there is an extensive history of work linking specific neuroanatomical properties with focused areas of cognition and behavior, only recently have large-scale collaborative efforts begun to provide the power necessary for data-driven discovery science (Somerville et al. 2018; Van Essen et al. 2013; Casey et al. 2018; Alexander et al. 2017; Holmes et al. 2015; Sudlow et al. 2015; Satterthwaite et al. 2014). Mounting evidence suggests that core features of brain anatomy are predictive of human behavior, but vary widely across populations and change across development within individuals (Bethlehem et al. 2021). Although this work has traditionally taken a cross-sectional, group-level approach, there is a growing understanding of the importance of accurately translating predictive models across both adult and developmental populations (Rosenberg, Casey, and Holmes 2018). To date, however, there is little empirical data on how the choice of both anatomical features and associated covariates may impact predictive accuracy or model generalizability, particularly within distinct demographic groups.

Across the cerebral cortex, individual variability in anatomical features, including surface area, gray matter volume, and cortical thickness, are predictive of a diverse set of behavioral traits, ranging from cognition (Seidlitz et al. 2018) to personality and mental health (Ooi et al. 2022). The establishment of meaningful imaging-based predictive models requires accurate and reliable measurements (Ge et al. 2017). Although, the within-sample predictions derived from brain anatomy generally account for less variance than those based on patterns of function and/or connectivity (Mansour et al. 2021; Dhamala, Jamison, Jaywant, Dennis, et al. 2021; Ooi et al. 2022), anatomical estimates are highly reliable (Holmes et al. 2015), highlighting their potential utility for brain-behavior predictive modeling. Relationships between neuroanatomical properties and cognitive abilities vary across the sexes (Gur et al. 1999; Gur and Gur 2016) and between healthy and clinical populations (Ehrlich et al. 2012; Hartberg et al. 2011) throughout the lifespan (Krogsrud et al. 2021). These studies, while crucial for the establishment of the neuroanatomical correlates of behavior, have largely focused on univariate analyses, leaving much to be understood about the multivariate associations that exist throughout the brain.

The volume of the cranium, typically referred to as intracranial volume (ICV) or estimated total intracranial volume, was historically thought to increase during development and remain stable throughout adulthood (Matsumae et al. 1996), with larger volumes in males than in females throughout the lifespan (De Bellis et al. 2001; Cosgrove, Mazure, and Staley 2007). More recent studies have confirmed greater ICV in males as well as changes in ICV throughout the lifespan (Caspi et al. 2020; Mills et al. 2016). ICV shows significant increases throughout childhood and adolescence (Mills et al. 2016; Dong, Castellanos, et al. 2020), followed by gradual increases in early adulthood until the fourth decade of life after which it begins to decrease (Caspi et al. 2020). These changes in ICV parallel shifts in cortical expansion (Hill et al. 2010), myelination (Grydeland et al. 2019), structure-function coupling (Baum et al. 2020), and functional maturation of association networks (Dong, Margulies, et al. 2020) throughout the lifespan (Sydnor et al. 2021). Across development, males exhibit larger ICV relative to females, along with a steeper rate of change during childhood and adolescence, as well as higher reduction rates after the fifth decade of life (Caspi et al. 2020). When investigating neuroanatomical properties (and their relationships to behavior) across the sexes and in different age groups, it is standard to correct for variations in ICV (Pintzka et al. 2015; Buckner et al. 2004), as corrected properties are assumed to be more valid than uncorrected measures (Sanfilipo et al. 2004), providing regional anatomical estimates unbiased by global shifts in head size across the population. Critically, ICV itself is also related to behavioral and psychological constructs of interests, including cognition (Van Loenhoud et al. 2018; MacLullich et al. 2002). Accordingly, the use of ICV correction may influence relationships between neuroanatomical properties and other variables of interest, an effect that could vary in impact across populations.

The presence of sex and/or gender differences in brain anatomy (e.g., Joel et al. 2015; Chekroud et al. 2016) and associated brain-behavior relationships has been a subject of pointed debate in the field (Ingalhalikar et al. 2014; Wierenga et al. 2019; Gur and Gur 2016). Several groups have investigated multivariate brain-behavior relationships in males and females (Dhamala, Jamison, Jaywant, and Kuceyeski 2021; Jiang, Calhoun, Cui, et al. 2020; Jiang, Calhoun, Fan, et al. 2020). While some data suggests the presence of dissociable functional connections and neuroanatomical features underlying cognitive abilities in men and women (Jiang, Calhoun, Cui, et al. 2020; Jiang, Calhoun, Fan, et al. 2020), other work has revealed that both sexes rely on shared functional connections (Dhamala, Jamison, Jaywant, and Kuceyeski 2021). However, this prior work has largely neglected the possible differential impact of ICV correction across the sexes, which could serve to amplify or mask group-level predictive features. Along these lines, recent work indicates that ICV correction methods can reduce both univariate sex differences and the accuracy of multivariate sex prediction based on gray matter volume (Sanchis-Segura et al. 2019; Sanchis-Segura et al. 2020). Meanwhile, in clinical populations, ICV correction can alter the relationships captured between regional neuroanatomical properties and behaviors (Voevodskaya et al. 2014). Although these data suggest ICV correction can influence both the results and their subsequent interpretation, it remains to be established whether these effects are consistent between sexes. Moreover, given the unique trajectories of ICV and regional neuroanatomy during development and adulthood (Caspi et al. 2020; Voevodskaya et al. 2014), it is likely that ICV correction will have differential effects across the lifespan.

In the current study, we sought to uncover the extent to which the most widely used ICV correction method, proportion-correction, might differentially bias brain-behavior predictions and associated interpretations across diverse populations. To directly address this open question, we investigated the sex-independent and sex-specific effects of accounting for individual differences in ICV using proportional corrections on predictions of cognition based on surface area, gray matter volume, and cortical thickness in healthy young adults from the Human Connectome Project (HCP) and typically developing children from the Adolescent Brain Cognitive Development (ABCD) dataset. First, examining the differences in predictive accuracy based on ICV-uncorrected and ICV-(proportion)-corrected neuroanatomical measures, we demonstrate that ICV correction reduces predictive accuracies achieved by sex-independent and sex-specific models based on surface area and gray matter volume but increases predictive accuracies achieved by models based on cortical thickness. Second, evaluating the effects of ICV correction on model generalizability across sexes and datasets, we determine that predictive models based on uncorrected measures of surface area and gray matter volume are more generalizable than their corrected counterparts. Conversely, ICV-corrected measures of cortical thickness are more generalizable than their uncorrected counterparts. Third, investigating the influence of ICV correction on the associations identified between neuroanatomical features and cognitive abilities, we reveal that distinct neuroanatomical features are associated with cognition across children and adults regardless of ICV correction, but those associations are shared across sexes for uncorrected measures of surface area and gray matter volume and for ICV-corrected measures of cortical thickness. Collectively, these results highlight the differential effects of ICV correction on behavioral predictions across neuroanatomical features, sexes, and age groups. Based on these findings, we speculate that ICV carries behaviorally relevant information, and this must be taken into consideration when developing predictive models to capture brain-behavior relationships across distinct populations that are likely to differ in ICV.

## Methods

An overview of our experimental workflow is shown in Figure 1. The methods used in this study build upon those previously described in our prior work (Dhamala, Jamison, Jaywant, Dennis, et al. 2021; Dhamala, Jamison, Jaywant, and Kuceyeski 2021; Anderson et al. 2021; Ooi et al. 2022) to perform novel analyses investigating effects of intracranial volume correction on sex-independent and sex-specific predictive modelling of behavioral traits in distinct populations.

**Figure 1:**
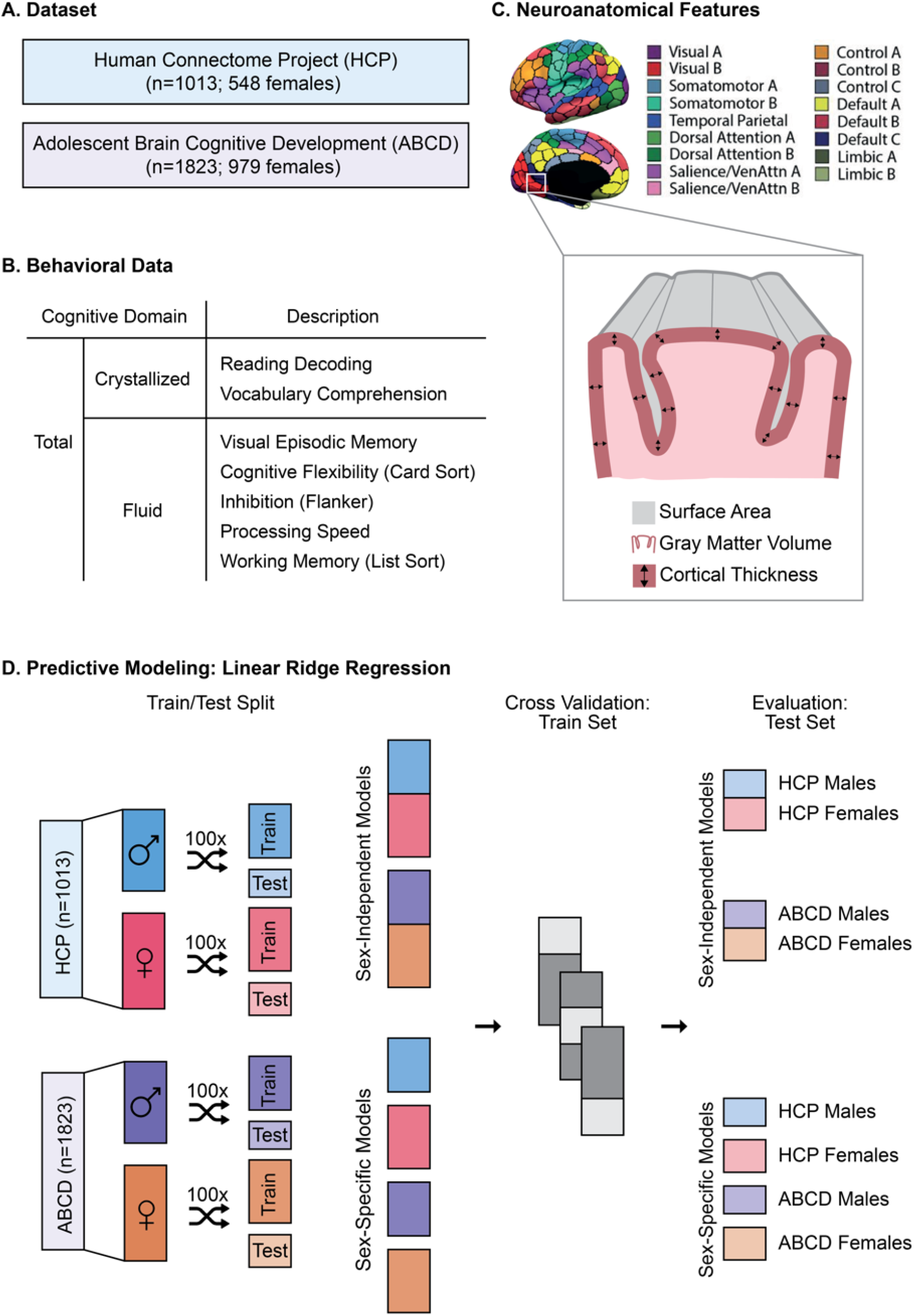
Experimental Workflow. (A) Dataset: Healthy young adults from the Human Connectome Project dataset, and typically developing children from the Adolescent Brain Cognitive Development dataset were included in the study. (B) Behavioral Data: Cognitive scores were compiled for each subject based on NIH Toolbox Cognitive Battery Task Scores. Total, Crystallized, and Fluid composites, as well as individual task scores within the Crystallized and Fluid domains were considered. (C) Neuroanatomical Features: Each subject’s native surface space was projected onto the 400-region Schaefer parcellation, and the T1-weighted anatomical image was used to extract regional surface area, gray matter volume, and cortical thickness for each of the 400 regions of interest. For each subject, these regional measures were either left uncorrected, or proportionally corrected for total intracranial volume. (D) Linear ridge regression models were trained to predict individual cognitive scores based on each of the neuroanatomical features. Males and females from each of the datasets were split into train (66%) and test sets. For each dataset, sex-independent models were trained on both male and female subjects, while sex-specific models were trained separately for each sex. All models employed three-fold cross-validation to optimize the regularization parameter. Sex-independent models were evaluated on sex-independent test sets from both datasets, and sex-specific models were evaluated on male- and female-specific test sets from both datasets.

### Datasets

We considered healthy young adult participants from the Human Connectome Project (HCP) – Young Adult S1200 release (Van Essen et al. 2013). The HCP dataset is a community-based sample of twins, siblings, and unrelated individuals who were assessed on a comprehensive set of neuroimaging and behavioral batteries. After pre-processing quality control of imaging data, as described in (Li et al. 2019; Kong et al. 2021; Ooi et al. 2022), we filtered participants based on availability of anatomical scans and behavioral scores of interests (Figure 1A). Our final HCP sample comprised 1013 adults (548 males; 22-37 years old). Although the term *gender* is used in the HCP Data Dictionary, the term *sex* is used in this article because the database collected self-reported biological sex information as opposed to gender identification. The self-reported biological sex information was not verified using genetic information.

We also considered typically developing children from the Adolescent Brain Cognitive Development (ABCD) 2.0.1 release (Casey et al. 2018). The ABCD dataset is a large community-based sample of children and adolescents who were assessed on a comprehensive set of neuroimaging, behavioral, and developmental batteries. After pre-processing quality control of imaging data, as described in (Chen et al. 2020; Ooi et al. 2022), including removal of participants from sites using Phillips scanners as recommended by the ABCD consortium, we filtered participants based on availability of anatomical scans and behavioral scores of interest (Figure 1B-C). Our final ABCD sample comprised 1823 children (979 males; 9-10 years old).

### Image Acquisition and Processing

Minimally processed T1-weighted anatomical images (0.7 mm isotropic for HCP; 1.0 mm isotropic for ABCD) were used for the analyses. Details about the acquisition protocol and processing pipelines are described elsewhere for HCP (Glasser et al. 2013; Marcus et al. 2013) and ABCD (Hagler et al. 2019).

### Behavioral Data

The NIH Toolbox Cognition Battery is an extensively validated battery of neuropsychological tasks used to assess language, executive function, episodic memory, processing speed, and working memory based on seven individual test instruments (Figure 1B) (Carlozzi et al. 2017; Gershon et al. 2013; Heaton et al. 2014; Mungas et al. 2014; Weintraub et al. 2013; Weintraub et al. 2014; Zelazo et al. 2014; Zelazo and Bauer 2013). Initial factor analysis of the individual task scores yields three composite scores: total, crystallized, and fluid. Broadly, crystallized cognition represents language abilities, while fluid cognition represents executive function, episodic memory, processing speed, and working memory. These composite scores tend to be more reliable and stable but may fail to capture variability in individual tasks (Heaton et al. 2014). Accordingly, all individual task scores as well as the composite scores were used in the analyses.

### Neuroanatomical Features

For each participant, the native *fs_LR32k* surface space was projected onto the 400-region Schaefer parcellation (Schaefer et al. 2018) using HCP workbench, and the T1-weighted anatomical image was used to extract cortical surface area, cortical gray matter volume, and cortical thickness for each of the 400 regions of interest (ROIs) using FreeSurfer 6.0’s *mris_anatomical_stats* (Dale, Fischl, and Sereno 1999) (Figure 1C). Measures of intracranial volume (ICV) were obtained from FreeSurfer’s estimated total intracranial volume. Surface area, gray matter volume, and thickness were proportionally corrected for individual differences in ICV by dividing the raw values by ICV. Both ICV-uncorrected and ICV-corrected anatomical measures were used in the analyses. Regional measures were also summarized at a network-level by computing the sum of the surface areas and gray matter volumes and the average of cortical thickness measures across all parcels within a network. Correlations between network-level measures of neuroanatomical features and ICV, as well as between ICV and the cognitive scores, were computed in a sex-specific manner for each dataset using Pearson’s correlation.

### Predictive Modelling

Sex-independent and sex-specific linear ridge regression models were trained to predict each behavioral score (individual task scores and composite scores) based on each anatomical measure (surface area, gray matter volume, thickness) (Figure 1D). Separate models were trained for ICV-uncorrected and ICV-corrected measures, as well as for each of the two datasets (HCP and ABCD). Sex-independent models were trained on data from all subjects, while male- and female-specific models were trained only on data from males and females, respectively. For each model, data were randomly shuffled and split into 100 distinct train (66%) and test sets without replacement. For the HCP data, family structure was considered when splitting the data such that related participants were placed either in the train or the test set but not split across both. Similarly, for the ABCD data, imaging site was considered such that all participants from a given site were placed either in the train or the test set but not split across both. Three-fold cross-validation was implemented to select and validate the regularization parameter using the train set. Family structure and imaging site were similarly accounted for in the cross-validation as in the initial train-test split. Once optimized, sex-independent models were evaluated on test sets from both datasets while sex-specific models were evaluated on test sets from both datasets and across both sexes. This was repeated for the 100 distinct train-test splits to obtain a distribution of performance metrics. The accuracy of each model is defined as the correlation between the true and predicted behavioral scores for each split. Average accuracy was computed by taking the mean across the 100 distinct train-test splits.

### Model Significance

All models were evaluated on whether they performed better than chance using null distributions of performance as previously described (Dhamala, Jamison, Jaywant, Dennis, et al. 2021; Dhamala, Jamison, Jaywant, and Kuceyeski 2021; Parkes et al. 2021). For each predictive model, the behavioral score was randomly permuted 10,000 times. Each permutation was used to train and test a null model using a randomly selected regularization parameter from the set of selected parameters for the original model. Prediction accuracy from each of the original model’s 100 train-test splits were then compared to the median prediction accuracy from the null distribution. The p-value for the model’s significance is defined as the proportion of 100 original models with prediction accuracies less than or equal to the median performance of the null model. In other words, a model is considered to be significant if it performed better than the median null performance for more than 95 of the 100 original models. The p values were then corrected for multiple comparisons across all cognitive scores using the Benjamini-Hochberg False Discovery Rate (q=0.05) procedure (Benjamini and Hochberg 1995).

### Model Comparisons

Models trained on ICV-uncorrected versus ICV-corrected anatomical measures were compared to one another to evaluate significant differences in performance using a two-tailed exact test of differences (MacKinnon 2009) as previously described (Dhamala, Jamison, Jaywant, Dennis, et al. 2021). The p-value for the model comparison is defined as the proportion of pairs of 100 models where one model’s prediction accuracy either is less than or equal to the other model. In other words, one model is considered to be significantly better than the other model if it performed better than the other for more than 95 of the 100 paired models. The p values were then corrected for multiple comparison across all cognitive scores using the Benjamini-Hochberg False Discovery Rate (q=0.05) procedure (Benjamini and Hochberg 1995).

### Model Generalizability

For sex-independent models, models trained on a given dataset were evaluated on both datasets. For each anatomical modality, an average model prediction accuracy was computed for each train/test dataset combination by taking the mean prediction accuracy across all cognitive scores. For sex-specific models, models trained on a given sex from a given dataset were evaluated on both sexes from both datasets. For each anatomical modality, a model prediction accuracy was computed for each train/test dataset and sex combination by taking the mean prediction accuracy across all cognitive scores. Model generalizability is defined as the accuracy obtained when a given model is evaluated on a population (i.e., a given sex and/or dataset) that is unique from the population in which it was trained. This is distinct from the model accuracy which is defined as the accuracy obtained when evaluating the model on a (hold out) test set that is from the same population as the training set. Model generalizability was also computed separately for each of the three cognitive domains; total (composite score only), crystallized (composite score and individual task scores), and fluid (composite score and individual task scores). Percent difference (% difference) between the ICV-uncorrected (r_uncorrected_) and ICV-corrected (r_corrected_) prediction accuracies was calculated as follows:

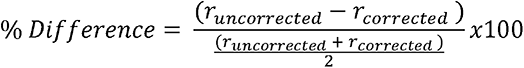

### Feature Weights

Raw feature weights obtained from the linear ridge regression models were transformed used the Haufe transformation (Haufe et al. 2014) to increase their interpretability and reliability (Tian and Zalesky 2021; Chen et al. 2020). For each train-test split, the raw feature weights, *W*, the covariance of the input data (anatomical modality) from that train set, Σ*_X_*, and the covariance of the output data (behavioral score) from that train set, Σ*_y_*, were used to compute the Haufe-transformed feature weights, *A*, as follows:

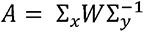

These Haufe-transformed feature weights were then averaged across the 100 splits to obtain a mean regional feature weight. To compare pairs of regression models, correlations between mean region feature weights were evaluated using Pearson’s correlation. Absolute regional feature weights were mapped to a network-level by assigning each Schaefer cortical parcel to one of 17 networks from the Yeo 17-network parcellation (Yeo et al. 2011) to generate network-level feature weights. Divergence in feature weights between models were evaluated using exact tests for differences.

### Data and Code Availability

All data used in this study are openly available and can be accessed directly from the HCP (https://www.humanconnectome.org/study/hcp-young-adult) and ABCD (https://abcdstudy.org/) websites. Code used to generate the results presented here are available on GitHub (https://github.com/elvisha/neuroanatomical-predictions-of-behaviour).

## Results

### Intracranial volume is uniquely related to brain anatomy across populations

Regional uncorrected and proportion-corrected measures of surface area, gray matter volume, and cortical thickness were summarized at the network-level by taking the sum of the surface areas and gray matter volumes and average of cortical thickness measures across all parcels within a network. These uncorrected and proportion-corrected network summaries of the anatomical properties were then correlated with total intracranial volume (ICV) to evaluate network-specific relationships between the neuroanatomical features and ICV, as shown in Figure 2.

**Figure 2:**
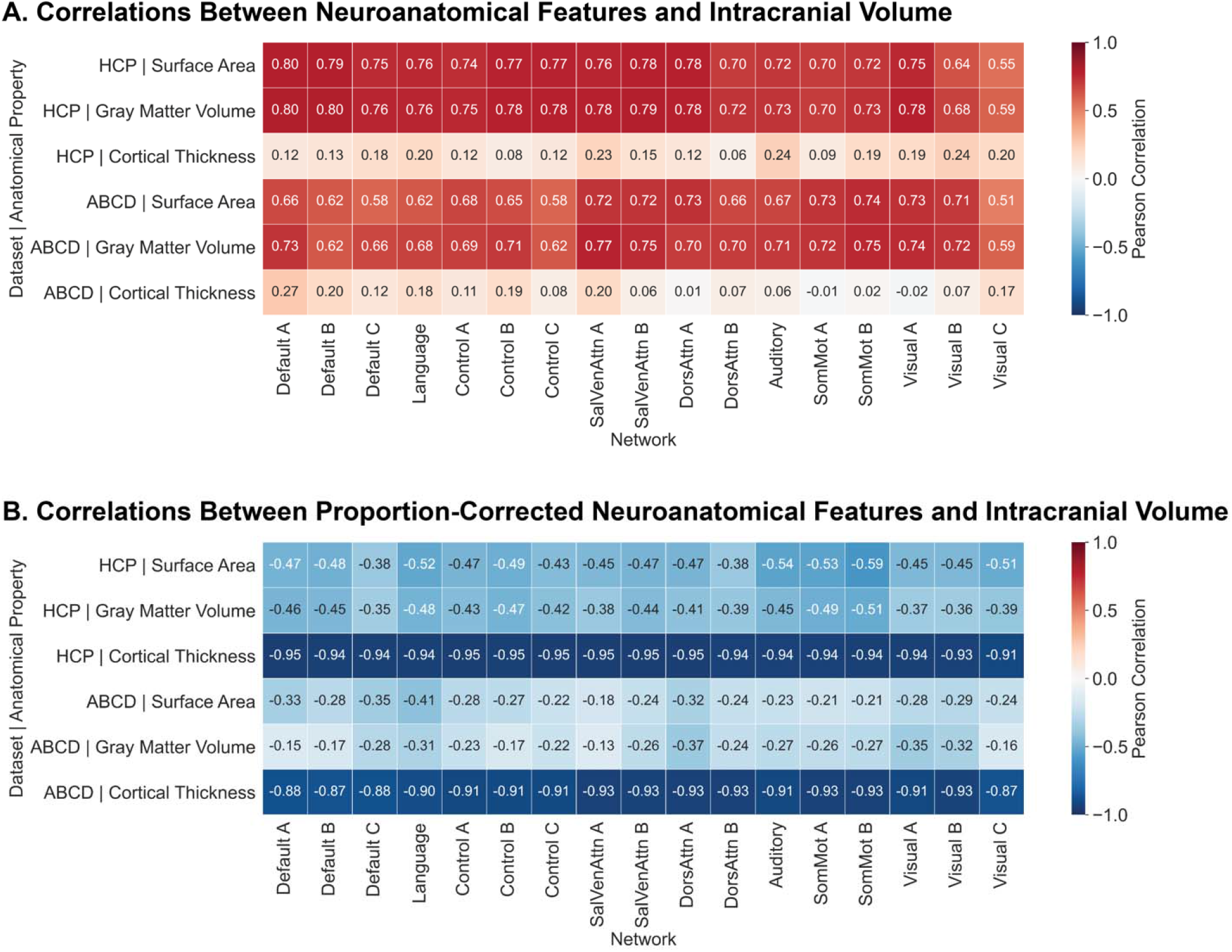
Total surface area and gray matter volume, and average cortical thickness across distinct cortical networks exhibit correlations with intracranial volume. Correlation (Pearson’s correlation coefficient) between network-level uncorrected (A) and proportion-corrected (B) neuroanatomical properties (total surface area, total gray matter volume, and average cortical thickness) and total intracranial volume across both datasets (HCP and ABCD). SalVenAttn – Salience/Ventral Attention; DorsAttn – Dorsal Attention; SomMot – Somatomotor. Networks are ordered from heteromodal (left) to unimodal (right).

Within the HCP dataset (Figure 2A), correlations between uncorrected surface area and ICV ranged between 0.55 and 0.80 (mean ± standard deviation=0.73 ± 0.06), with somewhat stronger relationships present in heteromodal association cortices (0.77 ± 0.02) than in unimodal somatosensory/motor (somato/motor) cortices (0.68 ± 0.07). Similarly, correlations between uncorrected gray matter volume and ICV ranged between 0.59 and 0.80 (0.75 ± 0.05) with somewhat stronger relationships in heteromodal association cortices (0.78 ± 0.02) than in unimodal cortices (0.70 ± 0.06). However, for uncorrected cortical thickness, correlations were generally weaker and ranged between 0.06 and 0.24 (0.16 ± 0.05), and somewhat stronger relationships were observed in unimodal (0.19 ± 0.05) than association cortices (0.14 ± 0.04).

Within the ABCD dataset (Figure 2A), correlations between ICV and uncorrected surface area or gray matter volume were generally comparable across unimodal and heteromodal cortices. Correlations between uncorrected surface area and ICV ranged between 0.51 and 0.74 (0.67 ± 0.06), while those between uncorrected gray matter volume and ICV ranged between 0.59 and 0.77 (0.69 ± 0.05). However, correlations between uncorrected cortical thickness and ICV exhibited the opposite pattern as that observed in HCP: correlations ranged between −0.02 and 0.30 (0.10 ± 0.08) with stronger relationships in heteromodal association cortices (0.14 ± 0.07) than in unimodal somato/motor cortices (0.05 ± 0.06). These data are consistent with prior work indicating a staggered maturation of cortical gray matter across development, in which the unimodal somato/motor and visual territories develop prior to the heteromodal association areas (Dong, Margulies, et al. 2020; Sydnor et al. 2021).

Across both datasets, correlations between all proportion-corrected neuroanatomical features and ICV were negative and comparable across unimodal and heteromodal cortices (Figure 2B). In HCP, the correlations ranged between −0.38 and −0.59 (−0.47 ± 0.05) for surface area, −0.35 and −0.51 (−0.43 ± 0.05) for gray matter volume, and −0.91 and −0.95 (−0.94 ± 0.01) for cortical thickness. In ABCD, they ranged between −0.18 and −0.41 (−0.27 ± 0.06) for surface area, −0.13 and −0.37 (−0.24 ± 0.07) for gray matter volume, and −0.87 and −0.93 (−0.91 ± 0.02) for cortical thickness. These data suggest that proportion-correction is unsuccessful in removing all of the variance related to individual differences in ICV for surface area and gray matter volume, and instead introduces additional information about ICV into cortical thickness measures.

Similar results were observed when correlations between ICV and the neuroanatomical properties were evaluated in a sex-specific manner in HCP (Figure S1A, S2A) and ABCD (Figure S1B, S2B).

These data suggest that ICV is differentially related to structural organization of the cortex during childhood and adulthood. Moreover, correcting for individual differences in ICV using the proportion-correction method can induce distinct effects in populations across the lifespan and inadvertently introduce information about intracranial volume into the neuroanatomical measures.

### Intracranial volume is distinctly related to abilities across cognitive domains

Sex-specific correlations between ICV and the ten cognitive scores (three composite, seven individual task) were computed (Figure S3). Across both datasets and sexes, correlations between ICV and the total and crystallized composite scores ranged between 0.16 and 0.24. Similar correlations ranging from 0.14 to 0.22 were observed with individual task scores within the crystallized domain. Generally weaker relationships were identified between ICV and the fluid composite (0.08 – 0.18) as well as individual fluid task scores (0.00 – 0.15) with the exception of the Working Memory (List Sort) score (0.13 – 0.19).

Relationships between brain volume and general intelligence have been previously reported (Gignac and Bates 2017). Here, in both children and adults, crystallized and fluid domains of cognition are differentially related to ICV, suggesting ICV may carry behaviorally relevant information that is partially distinct across cognitive domains.

### Intracranial volume correction differentially biases prediction accuracies across neuroanatomical features

Sex-independent models were trained to predict ten distinct behavioral scores using either ICV-uncorrected or ICV-corrected anatomical measures of surface area, gray matter volume, or cortical thickness in both HCP (adults) and ABCD (children). Prediction accuracies obtained by these models are shown in Figure 3.

**Figure 3:**
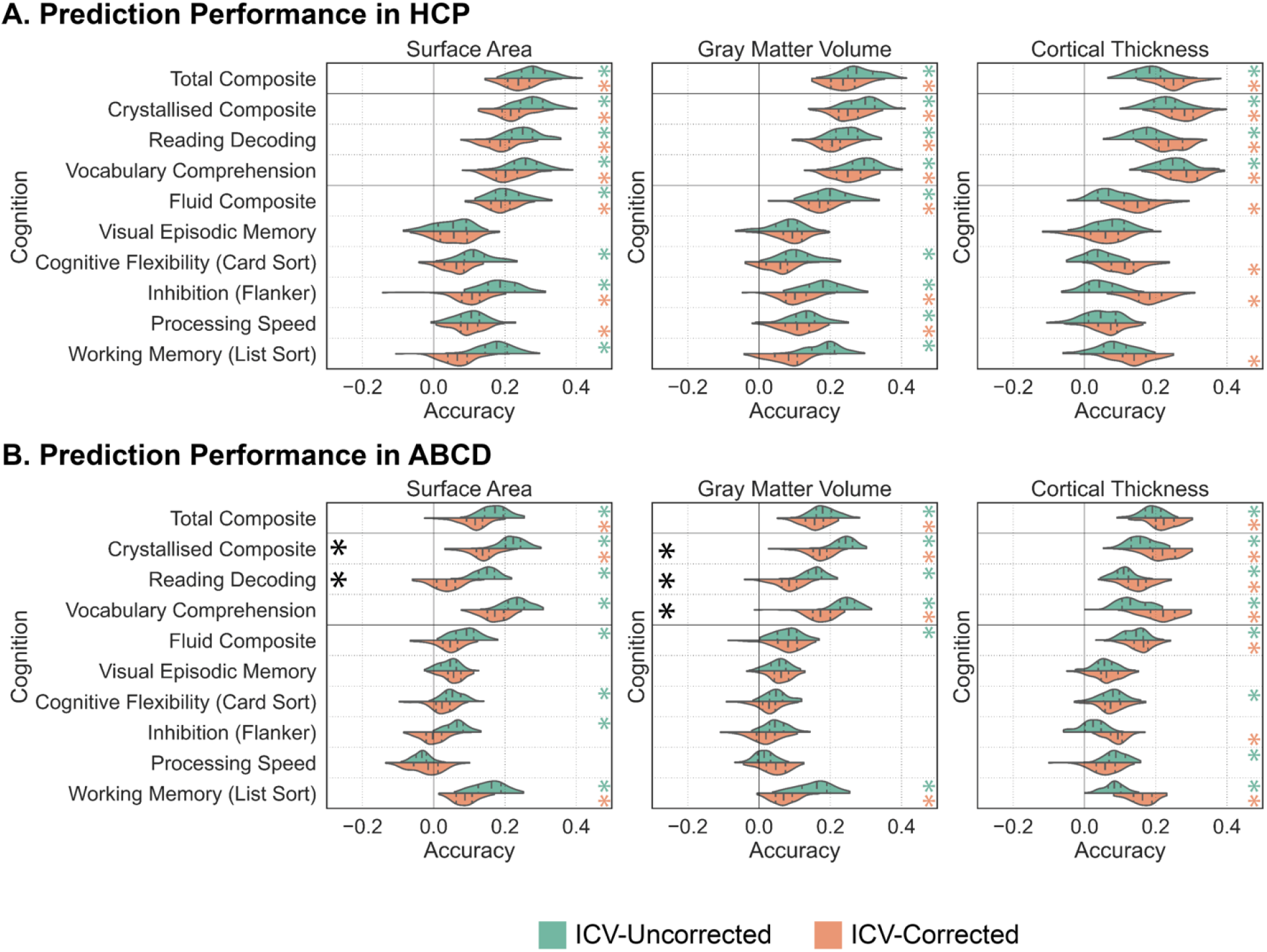
Accounting for intracranial volume reduces predictive accuracies of cognition based on surface area and gray matter volume, and increases predictive accuracies based on cortical thickness. Prediction accuracies (Pearson’s correlation coefficient between observed and predicted scores) for sex-independent models predicting cognitive scores in HCP (A) and ABCD (B). Predictions based on surface area (left), gray matter volume (middle), and cortical thickness (right) using ICV-uncorrected (green) and ICV-corrected (orange) anatomical properties are shown. Green and orange asterisks (*) denote that the model performed above chance levels based on permutation tests (corrected p<0.05). Black asterisks (*) denote that model performance was significantly different between the ICV-uncorrected and ICV-corrected predictions based on exact tests for differences (corrected p<0.05). The shape of the violin plots indicates the entire distribution of values, dashed lines indicate the median, and dotted lines indicate the interquartile range.

Sex-independent models based on uncorrected and corrected measures of surface area, gray matter volume, and cortical thickness successfully predicted (corrected p<0.05) total composite scores and scores within the crystallized domain in both datasets, but generally only successfully predicted scores within the fluid domain in HCP.

Models trained to predict cognitive scores in HCP achieved higher mean prediction accuracies based on uncorrected measures of surface area (r=0.190 for ICV-uncorrected, r=0.142 for ICV-corrected) and gray matter volume (r=0.200 for ICV-uncorrected, r=0.153 for ICV-corrected), and corrected measures of cortical thickness (r=0.119 for ICV-uncorrected, r=0.174 for ICV-corrected) (Figure 3A). Similarly, models trained to predict cognitive scores in ABCD yielded higher within-dataset mean prediction accuracies based on uncorrected measures of surface area (r=0.114 for ICV-uncorrected, r=0.064 for ICV-corrected) and gray matter volume (r=0.123 for ICV-uncorrected, r=0.087 for ICV-corrected), and corrected measures of cortical thickness (r=0.106 for ICV-uncorrected, r=0.142 for ICV-corrected) (Figure 3B). While this general trend was evident across analyses, most of these differences were non-significant at the level of the individual cognitive scores being predicted (corrected p>0.05). The one notable exception: models predicting cognitive scores within the crystallized domain in ABCD were significantly more accurate when using uncorrected measures of surface area or gray matter volume than when using their corrected counterparts.

Previous work has demonstrated that transformation of neurobiological variables can strengthen or weaken the brain-behavior associations captured by predictive models (Li et al. 2019). Our current findings show that ICV correction can similarly strengthen or weaken relationships between neuroanatomical features and individual cognitive abilities, revealing unique impacts in prediction accuracies across children and adults.

### Effects of intracranial volume correction on prediction accuracies differ across sexes and age groups

Sex-specific models were trained to predict ten distinct behavioral scores using either ICV-uncorrected or ICV-corrected anatomical measures of surface area, gray matter volume, or cortical thickness in both HCP and ABCD (Figure 4).

**Figure 4:**
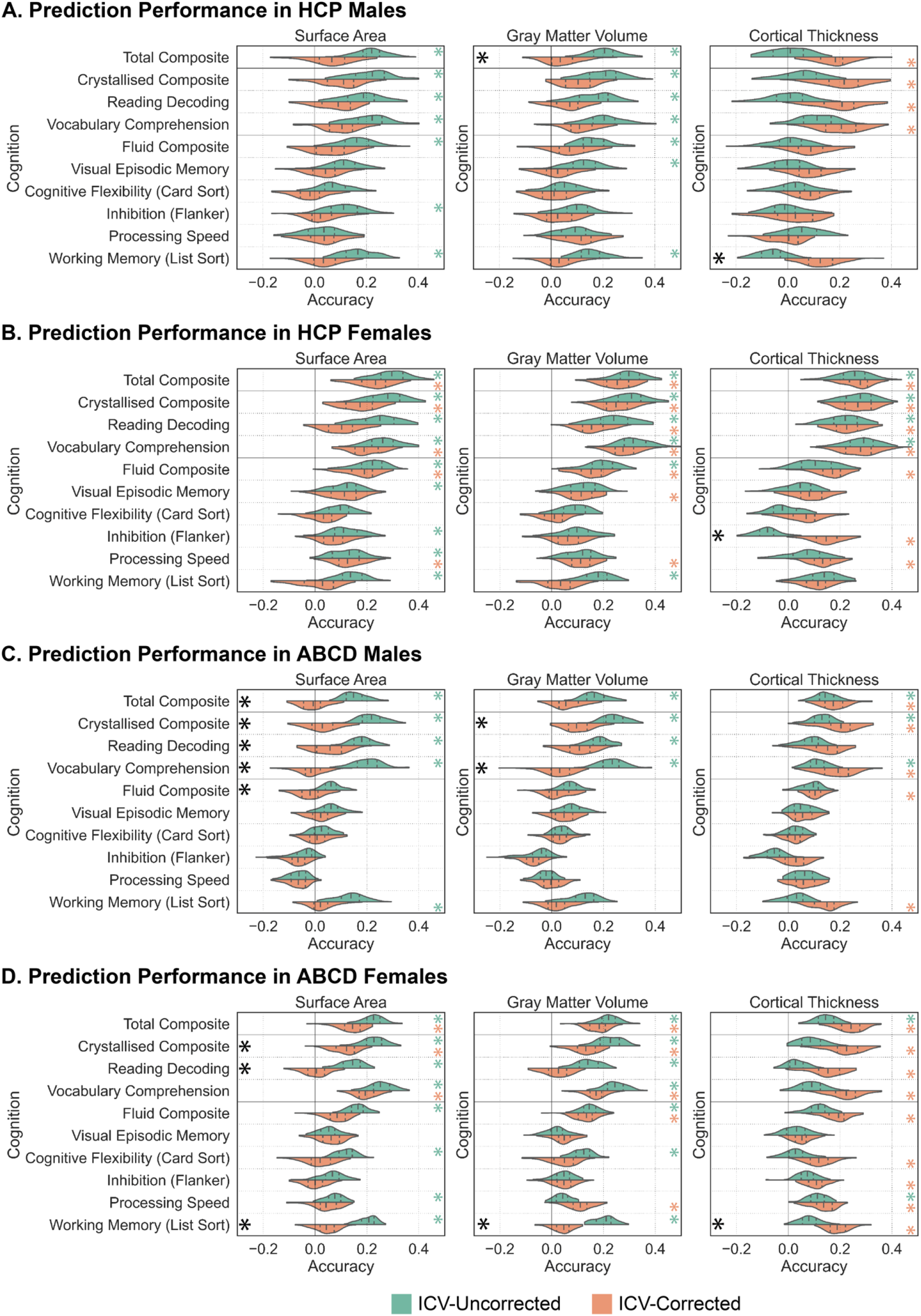
Accounting for intracranial volume differentially impacts predictive accuracies across surface area, gray matter volume, and cortical thickness in a sex specific manner. Prediction accuracies (Pearson’s correlation coefficient between observed and predicted scores) for sex-specific models predicting cognitive scores in HCP males (A), HCP females (B), ABCD males (C), and ABCD females (D). Predictions based on surface area (left), gray matter volume (middle), and cortical thickness (right) using ICV-uncorrected (green) and ICV-corrected (orange) anatomical properties are shown. Green and orange asterisks (*) denote that the model performed above chance levels based on permutation tests (corrected p<0.05). Black asterisks (*) denote that model performance was significantly different between the ICV-uncorrected and ICV-corrected predictions based on exact tests for differences (corrected p<0.05). The shape of the violin plots indicates the entire distribution of values, dashed lines indicate the median, and dotted lines indicate the interquartile range.

Within datasets, sex-specific models based on uncorrected and corrected measures of surface area, gray matter volume, and cortical thickness generally successfully predicted (corrected p<0.05) total composite scores and scores within the crystallized domain in females in both datasets. Sex-specific models generally successfully predicted (corrected p<0.05) total composite scores and scores within the crystallized domain in males within both datasets when using uncorrected measures of surface area or gray matter volume, or ICV-corrected measures of cortical thickness. Models trained to predict cognitive scores in HCP males achieved higher mean prediction accuracies when based on uncorrected measures of surface area (r=0.149 for ICV-uncorrected, r=0.053 for ICV-corrected) and gray matter volume (r=0.146 for ICV-uncorrected, r=0.061 for ICV-corrected), and corrected measures of cortical thickness (r=0.019 for ICV-uncorrected, r=0.119 for ICV-corrected) (Figure 4A). Similarly, models trained to predict cognitive scores in HCP females achieved higher mean prediction accuracies when based on uncorrected measures of surface area (r=0.188 for ICV-uncorrected, r=0.123 for ICV-corrected) and gray matter volume (r=0.194 for ICV-uncorrected, r=0.141 for ICV-corrected), and corrected measures of cortical thickness (r=0.128 for ICV-uncorrected, r=0.174 for ICV-corrected) (Figure 4B).

Models trained to predict cognitive scores in ABCD demonstrated similar trends. Higher mean prediction accuracies were achieved in males using models based on uncorrected measures of surface area (r=0.093 for ICV-uncorrected, r=-0.003 for ICV-corrected) and gray matter volume (r=0.106 for ICV-uncorrected, r=0.035 for ICV-corrected), and ICV-corrected measures of cortical thickness (r=0.066 for ICV-uncorrected, r=0.111 for ICV-corrected) (Figure 4C). Likewise, in females, higher mean prediction accuracies were achieved using models based on uncorrected measures of surface area (r=0.152 for ICV-uncorrected, r=0.071 for ICV-corrected) and gray matter volume (r=0.140 for ICV-uncorrected, r=0.092 for ICV-corrected), and ICV-corrected measures of cortical thickness (r=0.079 for ICV-uncorrected, r=0.162 for ICV-corrected) (Figure 4D). While differences based on uncorrected and corrected measures were generally non-significant at the level of individual cognitive scores, there were two noteworthy exceptions. First, models trained on uncorrected measures of surface area and gray mater volume in ABCD males and females significantly outperformed those trained on corrected measures (corrected p<0.05) to predict cognitive scores within the crystallized domain. Second, models trained in ABCD females achieved significantly higher prediction accuracies (corrected p<0.05) using uncorrected measures of surface area and gray matter volume, and ICV-corrected measures of cortical thickness to predict the Working Memory task score. Finally, within-dataset prediction accuracies were typically numerically higher in females than in males in HCP and ABCD.

In line with previous work, these data highlight the presence of differential brain-behavior predictive relationships across the sexes (Dhamala, Jamison, Jaywant, and Kuceyeski 2021; Jiang, Calhoun, Fan, et al. 2020; Jiang, Calhoun, Cui, et al. 2020). These results also emphasize the unique effects of ICV correction not just across age groups, but also across sexes within a given age group.

### Intracranial volume differentially influences generalizability of predictive models across neuroanatomical features, sexes, and age groups

Sex-independent models trained on each dataset were evaluated across both datasets. Mean prediction accuracies (across all cognitive scores) obtained by these models when evaluated within and between datasets are shown in Figure S4A. Prediction accuracies obtained for each cognitive score when evaluating the models between datasets are shown in Figure S5.

Sex-independent models trained to predict cognitive scores in HCP (Figures S4A, S5A) were more generalizable to ABCD based on uncorrected measures of surface area (r=0.099 for ICV-uncorrected, 0.007 for ICV-corrected) and gray matter volume (r=0.093 for ICV-uncorrected, r=0.013 for ICV-corrected), and corrected measures of cortical thickness (r=-0.044 for ICV-uncorrected, r=0.079 for ICV-corrected). Similarly, models trained to predict cognitive scores in ABCD (Figures S4A, S5B) were more generalizable to HCP based on uncorrected measures of surface area (r=0.144 for ICV-uncorrected, r=-0.023 for ICV-corrected) and gray matter volume (r=0.118 for ICV-uncorrected, r=-0.009 for ICV-corrected), and corrected measures of cortical thickness (r=-0.036 for ICV-uncorrected, r=0.147 for ICV-corrected).

Sex-specific models trained on each sex from each dataset were evaluated across both sexes and both datasets. Mean prediction accuracies obtained by these models when evaluated within and between sexes and datasets are shown in Figure S4B. Prediction accuracies obtained for each cognitive score when evaluating the models between datasets are shown in Figures S6-S9. In brief, sex-specific models exhibited similar overall trends in generalizability as those described for the sex-independent models above. Male- and female-specific models were more generalizable across sexes within datasets than they were across datasets when based on uncorrected measures of surface area or gray matter volume, or corrected measures of cortical thickness (Figure S4B). Moreover, sex-specific models based on ICV-corrected measures of surface area or gray matter volume, or uncorrected measures of cortical thickness generally achieved negative mean prediction accuracies when evaluated between datasets (Figure S4B).

Given the unique relationships between ICV and cognition across the different cognitive domains (total, crystallized, and fluid) outlined in Figure S3, we next examined each cognitive domain separately (see Figure S10 for the sex-independent models, and Figures S11-S13 for the sex-specific models). Sex-independent models trained to predict total cognition were more generalizable across datasets based on uncorrected measures of surface area (Figure S10A, left panel) and gray matter volume (Figure S10A, center panel), and corrected measures of cortical thickness (Figure S10A, right panel). Unsurprisingly, similar results were obtained for models trained to predict crystallized (Figure S10B) and fluid (Figure S10C) abilities, albeit at lower prediction accuracies for the fluid abilities.

Sex-specific models across each cognitive domain exhibited similar results as the sex-independent ones (see Figures S11-S13 for the sex-specific models). Models predicting total (Figure S11), crystallized (Figure S12), and fluid (Figure S13) abilities were more generalizable to the opposite sex within datasets than they were to either sex in the opposite dataset when based on uncorrected measures of surface area and gray matter volume, or corrected measures of cortical thickness. Models typically did not generalize between datasets (i.e., achieved negative prediction accuracies) when based on corrected measures of surface area and gray matter volume, or uncorrected measures of cortical thickness.

Percent differences in accuracy between models based on the ICV-uncorrected and ICV-corrected measures were also computed for each cognitive domain to quantify the effect of ICV correction and are shown in Figure 5A for the sex-independent models and Figure 5B for the sex-specific models. It is worth noting that in cases where one or both of the models being compared performed poorly, small absolute differences in accuracy can lead to exceptionally large percent differences. Hence, our results and interpretations focus on the general improvement/reduction of prediction accuracy rather than the numerical percent difference between models. As per the results described above, ICV correction reduced generalizability of models based on surface area (Figure 5A-B, left panels) and gray matter volume (Figure 5A-B, center panels), and increased generalizability of models based on cortical thickness (Figure 5A-B, right panels). Moreover, while accuracies were generally lower for predictions of fluid abilities, percent differences between the uncorrected and corrected measures are broadly comparable across the cognitive domains for sex-independent and sex-specific models. For sex-independent models, greater effects of ICV correction are observed in all three cognitive domains when evaluating model generalizability across sexes/datasets than model accuracy within a given sex and dataset. Models trained to predict total cognition exhibited lower absolute percent differences within datasets than between datasets for surface area (Figure 5A, top left panel), gray matter volume Figure 5A, top center panel), and cortical thickness (Figure 5A, top right panel). Similar patterns of larger effects of ICV correction for predictions across datasets than within datasets were observed for sex-independent predictions of crystallized (Figure 5A, middle panels) and fluid abilities (Figure 5B, bottom panels). Sex-specific models to predict total (Figure 5B, top panels), crystallized (Figure 5B, middle panels), and fluid (Figure 5B, bottom panels) abilities exhibited similar trends such that stronger effects of ICV correction were present when evaluating model predictions across datasets and sexes, than within datasets. Within datasets, predictions within and between sexes were generally similarly influenced by ICV correction in HCP and in ABCD.

**Figure 5:**
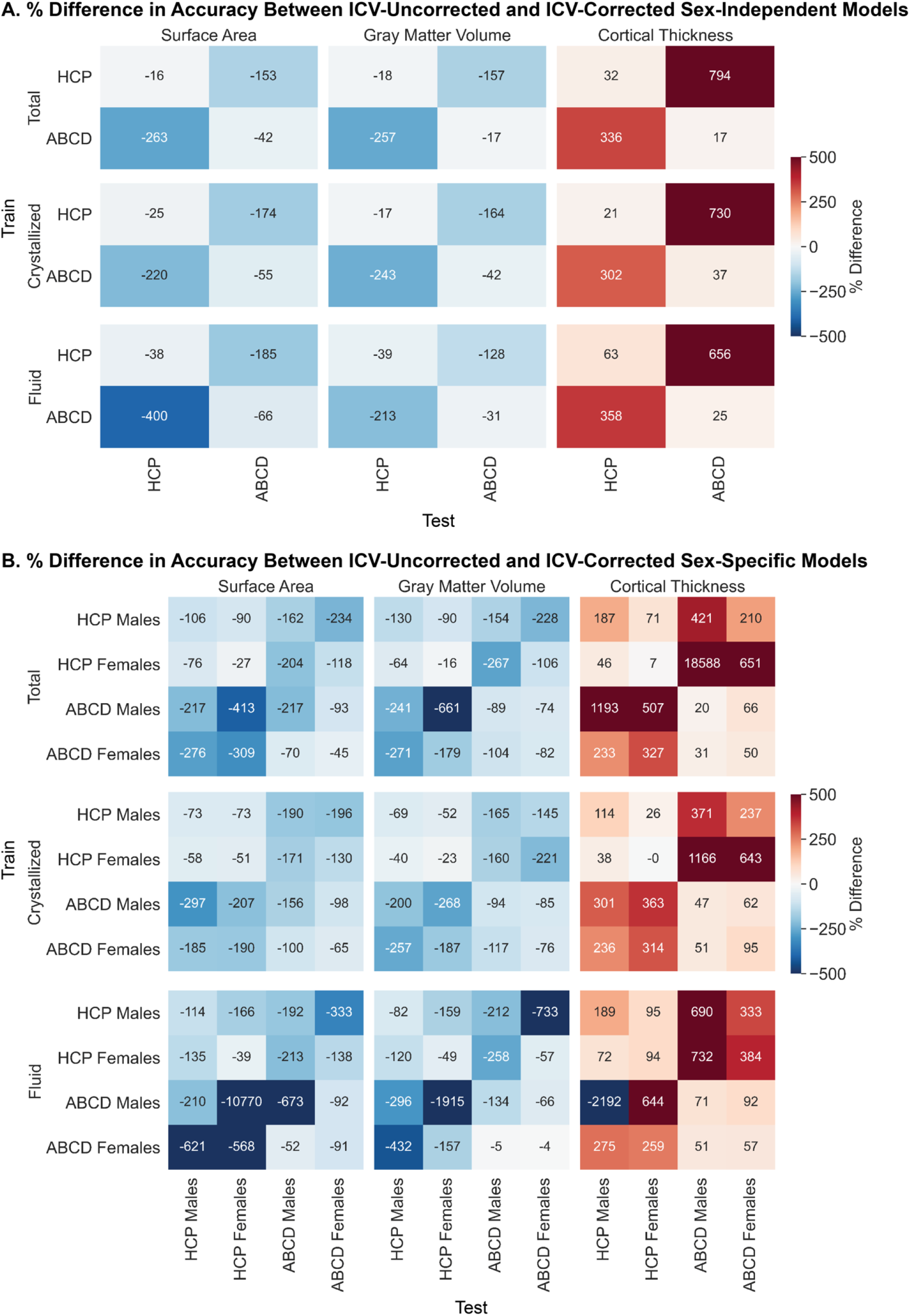
Intracranial volume correction differentially reduces generalizability of models based on surface area and gray matter volume but increases generalizability of models based on cortical thickness across cognitive domains. Percent difference in generalizability of sex-independent (A) and sex-specific (B) models across sexes (males and females) and datasets (HCP and ABCD). Percent difference between average prediction accuracies is based on ICV-uncorrected and ICV-corrected measures of surface area (left), gray matter volume (middle), and cortical thickness (right) to predict total (top), crystallized (middle), and fluid (bottom) abilities. Positive (warmer) values indicate that the ICV-corrected measures outperformed the ICV-uncorrected measures. The populations that the models were trained on are shown along the rows, and the populations that the models were tested on are shown along the columns.

Out-of-sample validation of predictive models using external datasets is typically considered the gold standard. These findings reveal that ICV correction more strongly affects out-of-sample predictions using external datasets than predictions on hold-out test sets within a dataset. These data also show that the correction affects males and females equally. Although speculative, this perhaps reflects differences in brain-behavior relationships throughout development and adulthood.

### Intracranial volume correction uniquely affects interpretations of brain-behavior relationships across neuroanatomical features and age groups

Regional Haufe-transformed feature weights were summarized at a network-level based on the Yeo 17-network solution (Yeo et al. 2011). Absolute relative network-level feature weights are shown in Figure 6 for the sex-independent models and in Figure S14-S15 for the sex-specific models. Within each dataset, measures of surface area and gray matter volume exhibit similar associations with individual cognitive abilities regardless of ICV correction. In HCP, surface area within the visual networks and gray matter volume within the default mode, language, and control networks are strongly associated with cognition (Figure 6A, left and center panels). In ABCD, surface area within the default mode network, and to a lesser extent the somatomotor network, as well as gray matter volume in the visual, somatomotor, and dorsal attention networks are strongly associated with cognition (Figure 6B, left and middle panels). Uncorrected measures of cortical thickness across a diverse set of networks are associated with cognitive abilities in both datasets (Figure 6A-B, top right panels). However, with ICV correction, opposing gradients of associations emerge in HCP and ABCD (Figure 6A-B, bottom right panels). In HCP, cortical thickness of regions within heteromodal association cortices are most strongly associated with cognitive scores while regions within unimodal cortices are weakly associated. However, in ABCD, this pattern is reversed and cortical thickness of regions within unimodal somato/motor cortices are most strongly associated with cognitive scores while regions within heteromodal association cortices are weakly associated.

**Figure 6:**
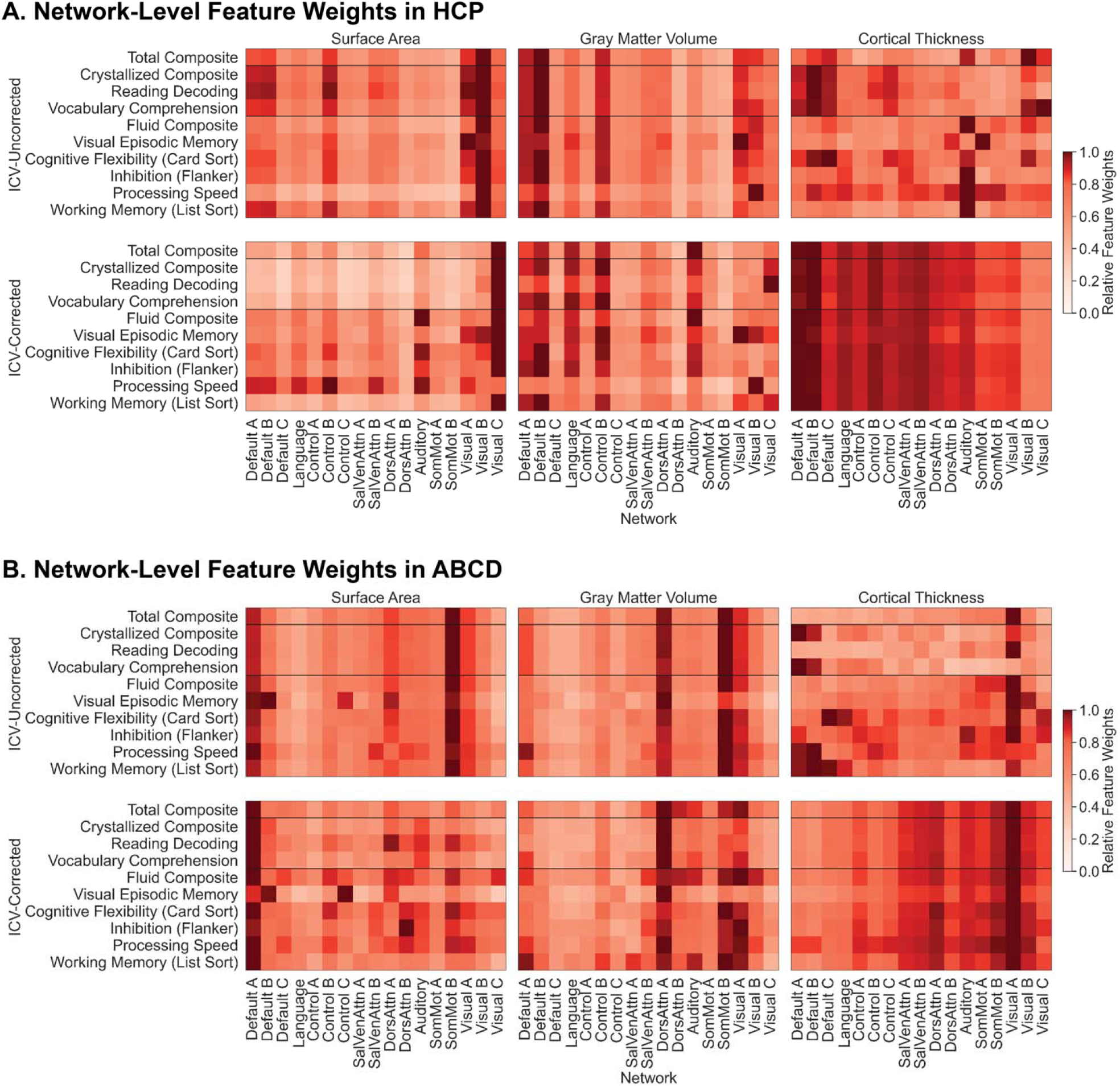
The predictive relationships linking cognition with the anatomy of association and unimodal cortices across populations can be revealed or obscured though the use of intracranial volume correction. Absolute relative network-level feature weights to predict each of the cognitive scores in HCP (A) and ABCD (B). Feature weights for models based on surface area (left), gray matter volume (middle), and cortical thickness (right) using ICV-uncorrected anatomical properties are shown in the top panels, and predictions using ICV-corrected anatomical properties are shown in the bottom panels. SalVenAttn – Salience/Ventral Attention; DorsAttn – Dorsal Attention; SomMot – Somatomotor. Networks are ordered from heteromodal (left) to unimodal (right).

These results emphasize that while ICV correction may not always affect network-level interpretations of brain-behavior relationships, it can reveal underlying relationships. The specific brain-behavior relationships we do capture, and the unique patterns they exhibit in the two datasets, are in line with prior work demonstrating non-linear maturation trajectories for cortical expansion, cortical thinning, intracortical myelination, functional maturation, and structure-function coupling (Sydnor et al. 2021) where unimodal somato/motor networks achieve maturity earlier in childhood followed by heteromodal association cortices later in adolescence.

### Intracranial volume correction differentially influences interpretations of brain-behavior relationships across sexes

Feature weights used to predict cognitive scores were extracted from the sex-specific models and Haufe-transformed. Regional surface area and gray matter volume feature weights to predict the Total Composite score are shown in Figure 7A, and regional cortical thickness feature weights to predict the Total Composite are shown in Figure 8A. Across sexes and datasets, there are widespread positive associations between the uncorrected measures of surface area and gray matter volume, and cognitive scores throughout the whole brain (Figure 7A). However, with ICV correction, HCP males and females demonstrated significantly weaker surface area (corrected p<0.05) and gray matter volume (corrected p<0.05) associations with cognition. While similar trends are present in ABCD males and females, the decrease in the strength of the associations is not significant. Regional feature weights of cortical thickness exhibit slightly different trends: in HCP, males and females exhibit diffuse positive and negative associations between uncorrected measures of cortical thickness and cognitive abilities, while in ABCD, males generally exhibit positive associations and females exhibit predominantly negative associations (Figure 8A). However, across both sexes and datasets, there exist widespread strong negative associations between ICV-corrected measures of cortical thickness and cognition. These differences in associations between the uncorrected and ICV-corrected measures were not significant.

**Figure 7:**
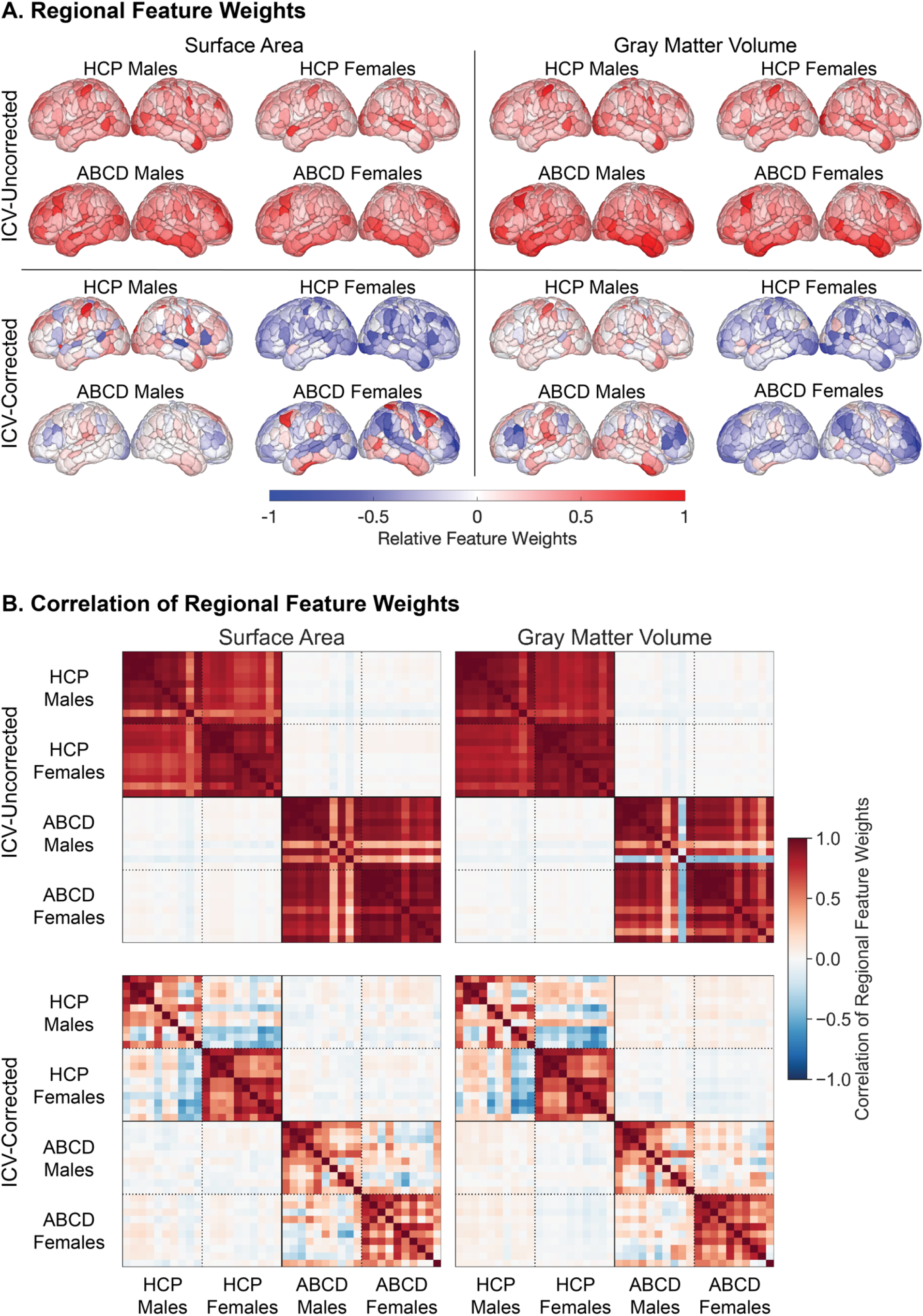
Regional surface area and gray matter volume associations with cognition are unique across age groups regardless of ICV correction, but they are shared across sexes without ICV correction and unique across sexes with ICV correction. Regional feature weights (A) and the correlations between them (B) for sex-specific models based on surface area (left) and gray matter volume (right). Models trained on HCP males, HCP females, ABCD males, and ABCD females using ICV-uncorrected anatomical properties are shown in the top panels, and predictions using ICV-corrected anatomical properties are shown in the bottom panels. Feature weights to predict total cognition are shown in (A) on lateral left (left) and right (right) cortical surfaces. Heatmaps of correlations of regional feature weights are ordered along the rows and columns based on the populations the models were trained on with in the following order: HCP Males, HCP Females, ABCD Males, ABCD Females. Within the blocks for each of those training sets, regional feature weights are ordered based on the cognitive scores being predicted as follows: Total Composite, Crystallised Composite, Reading Decoding, Vocabulary Comprehension, Fluid Composite, Visual Episodic Memory, Cognitive Flexibility (Card Sort), Inhibition (Flanker), Processing Speed, Working Memory (List Sorting).

**Figure 8:**
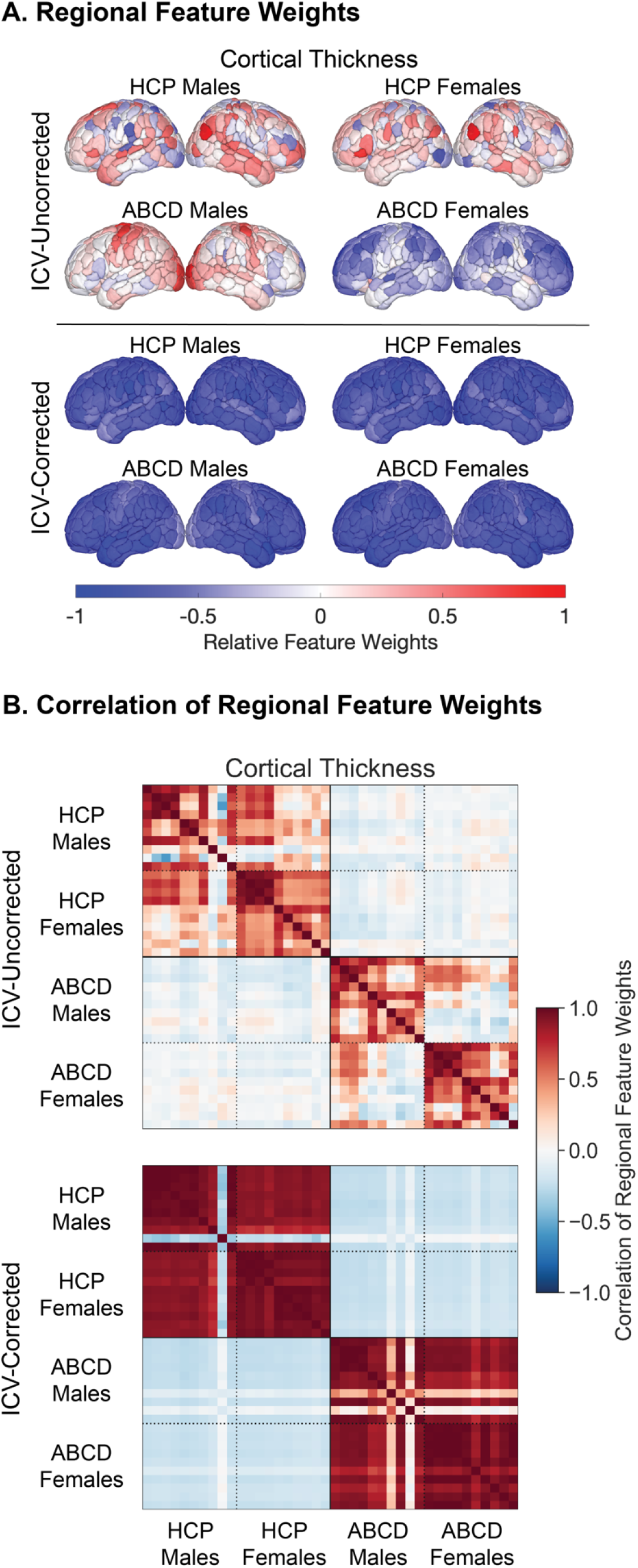
Associations between regional cortical thickness and cognition are unique across age groups regardless of ICV correction, but they are unique across sexes without ICV correction and shared across sexes with ICV correction. Regional feature weights (A) and the correlations between them (B) for sex-specific models based on cortical thickness. Models trained on HCP males, HCP females, ABCD males, and ABCD females using ICV-uncorrected anatomical properties are shown in the top panels, and predictions using ICV-corrected anatomical properties are shown in the bottom panels. Feature weights to predict total cognition are shown in (A) on lateral left (left) and right (right) cortical surfaces. Heatmaps of correlations of regional feature weights are ordered along the rows and columns based on the populations the models were trained on with in the following order: HCP Males, HCP Females, ABCD Males, ABCD Females. Within the blocks for each of those training sets, regional feature weights are ordered based on the cognitive scores being predicted as follows: Total Composite, Crystallised Composite, Reading Decoding, Vocabulary Comprehension, Fluid Composite, Visual Episodic Memory, Cognitive Flexibility (Card Sort), Inhibition (Flanker), Processing Speed, Working Memory (List Sorting).

Correlations between the feature weights were also analyzed and are shown for surface area and gray matter volume in Figure 7B, and for cortical thickness weights in Figure 8B. Across datasets, there is little to no overlap in the features used to predict cognitive abilities based on uncorrected measures of surface area (average correlation between regional feature weights, r=0.00) and gray matter volumes (r=-0.01) (Figure 7B, top panels). However, within datasets, male- and female-specific models rely on shared features to predict cognitive scores based on uncorrected measures of surface area (r=0.81 for HCP, r=0.81 for ABCD) and gray matter volume (r=0.86 for HCP, r=0.67 for ABCD). With ICV correction, across datasets, there remain little to no overlap in the features used by models based on surface area (r=0.00) or gray matter volume (r=0.01), but correlations observed within datasets between sexes are also generally reduced for both surface area (r=0.24 for HCP, r=0.29 for ABCD) and gray matter volume (r=0.24 for HCP, r=0.31 for ABCD) (Figure 7B, bottom panels). The opposite pattern is observed with cortical thickness: feature weights extracted from models using ICV-corrected measures are correlated across sexes (r=0.80 for HCP, r=0.78 for ABCD) but not datasets (r=-0.20), while those extracted from models using uncorrected measures are reduced across sexes (r=0.43 for HCP, r=0.32 for ABCD) and datasets (r=-0.04) (Figure 8B).

Correcting for ICV can reduce univariate sex differences in neuroanatomical properties as well as multivariate predictions of biological sex (Sanchis-Segura et al. 2019; Sanchis-Segura et al. 2020). Relationships between regional neuroanatomical properties and behaviors can also be altered by ICV correction in clinical populations (Voevodskaya et al. 2014). Our findings provide additional evidence that ICV correction can influence interpretation of regional-level brain-behavior relationships, particularly between sexes within a dataset. Based on these data, we emphasize that the unique effects of ICV correction across populations throughout the lifespan bias not only the strength of the brain-behavior relationships we can capture, but also their interpretability.

## Discussion

The application of predictive modelling in neuroimaging has provided foundational insights into the neurobiological correlates of behavior. While population-level associations between neuroanatomy and cognition have been extensively studied, prior predictive modeling work has not explicitly addressed the extent to which these relationships are shared across sexes and age groups. A standard practice when studying brain anatomy is to correct for individual differences in ICV using proportion correction, but the impact of this correction on the brain-behavior predictions had not previously been examined. Here, we demonstrate that proportional ICV correction differentially biases behavioral predictions and the subsequent interpretations of the underlying brain-behavior relationships across neuroanatomical properties, sexes, and age groups. For both the ABCD (n=1823; children) and HCP (n=1003; adults) datasets, the size of individual cortical regions (in terms of surface area and gray matter volume) predicts behavioral traits within and between sexes and datasets with greater accuracy and generalizability when ICV correction is not implemented. The captured associations between behavioral traits and regional surface area or gray matter volume are unique across children and adults regardless of ICV correction, but unique across sexes with ICV correction and shared otherwise. Conversely, regional cortical thickness predicts behavioral traits with greater accuracy and generalizability when individual differences in ICV are corrected. The associations between behavioral traits and regional cortical thickness are also consistently unique across children and adults but shared across sexes with ICV correction and unique otherwise. Taken together, these results reveal the differential effects of ICV correction on accuracy, generalizability, and interpretability of behavioral predictions across neuroanatomical features, sexes, and age groups.

There are marked differences in head size between individuals, presenting a challenge for the measurement of regional volumes and surface areas. Established differences in gray matter volume and ICV between the sexes and throughout the lifespan (Caspi et al. 2020; De Bellis et al. 2001; Bethlehem et al. 2021) have led to widespread implementations of ICV correction when studying brain-behavior relationships across populations. In this present work, we described how proportional ICV correction influences accuracy of behavioral predictions within a population, generalizability of the predictions across populations, and interpretations of the underlying brain-behavior relationships. We began by quantifying how ICV relates to uncorrected neuroanatomical properties and cognitive domains. Here, we observed diverging gradients in the network-specific relationships with uncorrected neuroanatomical properties across adults and children. In adults, surface area and gray matter volume across all networks were strongly related to ICV, but exhibited stronger correlations in heteromodal association cortices than in unimodal somato/motor cortices. Cortical thickness was weakly related to ICV across all networks, but exhibited stronger correlations with ICV in unimodal somato/motor cortices. In children, relationships with ICV were equally strong and comparable between unimodal somato/motor cortices and heteromodal association cortices for surface area and gray matter volume, while correlations with cortical thickness were weak overall but slightly stronger in heteromodal association cortices than unimodal cortices. These data are consistent with extensive work in neurodevelopment establishing that unimodal cortices exhibit earlier cortical expansion and cortical thinning than heteromodal cortices (Sydnor et al. 2021). We also quantified relationships between ICV and proportion-corrected neuroanatomical properties. In doing so, we observed that across both datasets and all networks, the relationships with surface area and gray matter volume were inverted and reduced in magnitude. Meanwhile, correlations between ICV and proportion-corrected cortical thickness were strongly negatively. We also noted that in adults and children, ICV is more strongly correlated with crystallized abilities than fluid abilities, which is in agreement with existing work (Farias et al. 2012). Collectively, these data reveal that shared relationships between ICV and cognitive domains exist during childhood and adulthood, but relationships between ICV and neuroanatomical features are unique across those populations. The analyses also demonstrate that proportional ICV correction does not entirely remove information pertaining to ICV from measures of surface area and gray matter volume, and actually introduces information about ICV to measures of cortical thickness. Therefore, this field standard method may not be ideal when accounting for individual differences in brain volume.

The rapidly growing use of predictive modelling in neuroimaging to map brain-behavior relationships has yielded numerous important advances in recent years. Studies have investigated how preprocessing (Li et al. 2019), data transformation (Parkes et al. 2021), predictive algorithms (He, Kong, et al. 2020), neuroimaging features (Dhamala, Jamison, Jaywant, Dennis, et al. 2021; Greene et al. 2018), model translation (He, An, et al. 2020), parcellation choices (Dhamala, Jamison, Jaywant, Dennis, et al. 2021), sample sizes (Marek et al. 2020), and phenotype selection (Chen et al. 2020) can influence neuroimaging-based predictions of individual behaviors. Unfortunately, these studies have in large part relied on single datasets of healthy young adults to train and evaluate model performance, even though it is becoming increasingly evident that models must be not only replicable and reliable within a dataset (Tian and Zalesky 2021), but also generalizable across datasets (Scheinost et al. 2019).

In this study, we quantified the extent to which predictive models generalize across distinct populations, evaluating whether this generalizability is influenced by ICV correction. Upon confirming how ICV is related to neuroanatomical features and cognition, we sought to determine how ICV correction differentially influences predictive models of behaviors based on the size or thickness of cortical territories in adults and children. ICV correction reduced (within sex and/or dataset) accuracy and (between sex and/or dataset) generalizability of predictions based on surface area and gray matter, and increased accuracy and generalizability based on cortical thickness. We speculate that these effects are driven by the inherent relationships that ICV has with both regional neuroanatomy and behavior. Given that surface area and gray matter volume are more strongly correlated to ICV than cortical thickness, and the proportion-correction reduces and inverts those relationships, ICV correction for measures of surface area and gray matter volumes results in the removal of behaviorally relevant information (captured in ICV) thus impairing predictions. Conversely, proportional ICV correction for cortical thickness introduces ICV-relevant information into the measures that may indirectly enhance the predictions. These observed effects were more pronounced in predictions across datasets than within datasets. As previously mentioned, ICV tends to differ between children and adults and these group differences may explain why models are more influenced by ICV correction when evaluating their generalizability across populations than within populations. Finally, these effects of ICV correction were surprisingly comparable across cognitive domains even though the cognitive domains themselves are differentially related to ICV. This suggests that the effect of ICV correction on prediction of a behavioral trait is, to some extent, independent of the relationship that ICV shares with that behavioral trait. Therefore, even in the absence of the underlying relationships between ICV and the behavior of interest, researchers must be aware of the influence that the correction may impart on their predictive modelling and subsequent interpretations. This work provides the basis for further exploration into whether ICV correction equally influences predictions of non-cognitive behaviors, including personality, and mental health.

The use of open-access neuroimaging datasets has gained considerable popularity in recent years (Madan 2021; Bzdok and Yeo 2017). Several studies have used these large-scale datasets to model brain-behavior relationships, but if and how those models can be interpreted is still up for debate (Tian and Zalesky 2021; Kohoutová et al. 2020). Here, we evaluated how ICV correction influences model interpretations of the neurobiological features that underlie individual cognitive abilities at both a network-level and a regional-level. At a network-level, feature weights extracted from models based on surface area and gray matter volume were generally unchanged with ICV correction. However, network-level features of cortical thickness demonstrated no interpretable trends without ICV correction, but a definitive gradient of network contributions emerged with ICV correction. These network-level weights across all neuroanatomical features were generally shared across sexes and unique across age groups regardless of ICV correction. At a regional level, examining the feature weights from the uncorrected models may lead one to conclude that relationships between the neuroanatomical features and cognition are unique across age groups but shared across sexes within age groups for surface area and gray matter volume but unique across sexes for cortical thickness. However, if ICV correction is implemented, we observe a different pattern: relationships remain unique across age groups, and are also unique across sexes within age groups for surface area and gray matter volume, but are shared across sexes for cortical thickness. Previously, we suggested that ICV correction inadvertently removes behaviorally relevant information from models based on surface area and gray matter volume and introduces it into models based on cortical thickness. The same mechanism may also explain the discrepancies in feature weights between uncorrected and ICV-corrected models. At the population-level males and females differ in ICV. However, this broad trend masks the presence of substantial variability and overlapping phenotypic distributions across populations. Predictions based on uncorrected measures of surface area and gray matter volume and corrected measures of cortical thickness may rely heavily on overlapping ICV information shared across the sexes (within each age group) while those based on corrected measures of surface area and gray matter volume and uncorrected measures are more reliant on unique relationships that are not driven by ICV. These findings serve as a cautionary tale for researchers using predictive modelling approaches to identify complex multivariate brain-behavior relationships across healthy and clinical populations without considering how factors such as ICV correction may be undermining their efforts and unintentionally biasing their interpretations.

Although many researchers have studied age- and sex-related differences in cortical organization and cognitive abilities, most prior experiments have focused on one or the other, or relied on univariate analyses (Cummings et al. 2020; Giedd and Rapoport 2010; Gong, He, and Evans 2011; Gur and Gur 2017; Jäncke 2018; Lenroot and Giedd 2010; Scheinost et al. 2015; Hagmann et al. 2010; Fair et al. 2009; Satterthwaite et al. 2015). Leveraging two large, open-access datasets, we quantified age-and sex-specific neurobiological correlates of cognition using multivariate predictive modelling approaches. Surface area of unimodal somato/motor regions, and gray matter volume and cortical thickness of heteromodal association regions were most strongly associated of cognition in adults. In children, the opposite was observed: surface area of heteromodal association regions, and gray matter volume and cortical thickness in unimodal somato/motor regions exhibited the strongest associations with cognition. Cortical surface area increases during childhood, before reaching a global peak at around 9 years of age and then slowly declining (Wierenga et al. 2014). Gray matter volume exhibits a similar trajectory but peaks occur between ages 11 and 14 (Gogtay and Thompson 2010). Global cortical thickness increases from birth until early childhood (Wang et al. 2019), before declining throughout adolescence and adulthood (Zhou et al. 2015). Studies of surface area, gray matter volume, and cortical thickness have established a progression of cortical maturation along the somatomotor-association axis: progression begins in unimodal somato/motor cortices and ends in heteromodal association cortices (Sydnor et al. 2021; Gogtay and Thompson 2010). The differential relationships we observe between the neuroanatomical properties and individual cognitive abilities in children and adults, suggest that brain-behavior relationships likely exhibit a similar developmental trajectory along the somatomotor-association axis as the neuroanatomical properties themselves. Given the cross-sectional nature of this study, we are limited in our ability to draw conclusions about these trajectories but future analyses incorporating longitudinal samples from ABCD will be able to capture them more definitively.

Of note, the findings of this study are subject to several limitations. First, these analyses are focused only on evaluating the effects the most widely used ICV correction method: the proportion correction. Other methods for the correction include covariate regression, non-linear modulation based on voxel-based morphometry (Good et al. 2001), the power-corrected proportion (Liu et al. 2014), and the residuals adjustment (Arndt et al. 1991; Mathalon et al. 1993). Methods that rely on population-level information (i.e., covariate regression, power-corrected proportion, and residuals adjustment) for the correction must be implemented separately within each cross-validation fold for every train-test split to prevent data leakage and then applied to the test set. Consequently, the model would have less utility when generalizing it to a population that is distinct from the training set (i.e., a different sex or age group). Second, the two datasets we relied on for this study capture a relatively small range of ages. HCP includes subjects between the ages of 22 and 37 while ABCD includes subjects who are 9-10 years old. Given this limited age range, we are unable to identify the network-level trajectories of brain-behavior relationships that exist throughout the lifespan. Future analyses of these relationships in adolescents and older adults can be used to supplement our findings and establish trajectories of associations between cognition and neuroanatomical organization in unimodal somatosensory and heteromodal association cortices. Finally, we rely on a single dataset for each age group studied, but the adults and children in these datasets are not necessarily representative of those populations. We cannot rule out the possibility that the effects observed here are driven by differences in imaging parameters or scanning acquisitions unrelated to the age differences. However, given that our findings align with established developmental cortical maturation trajectories (Blakemore 2012; Stiles and Jernigan 2010; Casey et al. 2005), it is likely that our results are capturing core age-related effects.

## Conclusion

An understanding the effects of data transformation on predictive models of brain-behavior relationships can enable to develop more accurate, generalizable, and interpretable models. In this work, we establish the differential impact of ICV correction on models of cognition based on cortical size and thickness in adults and children. Accuracy and generalizability were reduced with ICV correction for models based on size (surface area and gray matter volume) but increased for models based on thickness. Interpretability of the features that these models relied on were also affected by ICV correction: brain-behavior associations were unique across children and adults regardless of the correction, but only unique across sexes for models based on ICV-corrected measures of cortical size and uncorrected measures of cortical thickness. Taken together, these findings emphasize that we must carefully consider individual differences in ICV when evaluating brain-behavior associations across populations as those differences may influence the strength and associated interpretability of the underlying relationships.

## Acknowledgements

This work was supported by the National Institute of Mental Health (R01MH120080 and R01MH123245 to A.J.H.) and the Kavli Institute for Neuroscience at Yale University (Postdoctoral Fellowship for Academic Diversity to E.D.). This work was also supported by the following awards to B.T.T.Y.: the Singapore National Research Foundation (NRF) Fellowship (Class of 2017), the NUS Yong Loo Lin School of Medicine (NUHSRO/2020/124/TMR/LOA), and the Singapore National Medical Research Council (NMRC) LCG (OFLCG19May-0035).

Data were provided in part by the Human Connectome Project, WU-Minn Consortium (Principal Investigators: David Van Essen and Kamil Ugurbil; 1U54MH091657) funded by the 16 NIH Institutes and Centers that support the NIH Blueprint for Neuroscience Research; and by the McDonnell Center for Systems Neuroscience at Washington University.

Data used in the preparation of this article were also obtained from the Adolescent Brain Cognitive Development ^SM^ (ABCD) Study (https://abcdstudy.org), held in the NIMH Data Archive (NDA). This is a multisite, longitudinal study designed to recruit more than 10,000 children aged 9-10 and follow them over 10 years into early adulthood. The ABCD Study® is supported by the National Institutes of Health and additional federal partners under award numbers U01DA041048, U01DA050989, U01DA051016, U01DA041022, U01DA051018, U01DA051037, U01DA050987, U01DA041174, U01DA041106, U01DA041117, U01DA041028, U01DA041134, U01DA050988, U01DA051039, U01DA041156, U01DA041025, U01DA041120, U01DA051038, U01DA041148, U01DA041093, U01DA041089, U24DA041123, U24DA041147. A full list of supporters is available at https://abcdstudy.org/federal-partners.html. A listing of participating sites and a complete listing of the study investigators can be found at https://abcdstudy.org/consortium_members/. ABCD consortium investigators designed and implemented the study and/or provided data but did not necessarily participate in the analysis or writing of this report. This manuscript reflects the views of the authors and may not reflect the opinions or views of the NIH or ABCD consortium investigators.

## References

1. Alexander, L. M., J. Escalera, L. Ai, C. Andreotti, K. Febre, A. Mangone, N. Vega-Potler, N. Langer, A. Alexander, M. Kovacs, S. Litke, B. O’Hagan, J. Andersen, B. Bronstein, A. Bui, M. Bushey, H. Butler, V. Castagna, N. Camacho, E. Chan, D. Citera, J. Clucas, S. Cohen, S. Dufek, M. Eaves, B. Fradera, J. Gardner, N. Grant-Villegas, G. Green, C. Gregory, E. Hart, S. Harris, M. Horton, D. Kahn, K. Kabotyanski, B. Karmel, S. P. Kelly, K. Kleinman, B. Koo, E. Kramer, E. Lennon, C. Lord, G. Mantello, A. Margolis, K. R. Merikangas, J. Milham, G. Minniti, R. Neuhaus, A. Levine, Y. Osman, L. C. Parra, K. R. Pugh, A. Racanello, A. Restrepo, T. Saltzman, B. Septimus, R. Tobe, R. Waltz, A. Williams, A. Yeo, F. X. Castellanos, A. Klein, T. Paus, B. L. Leventhal, R. C. Craddock, H. S. Koplewicz, and M. P. Milham. 2017. ‘An open resource for transdiagnostic research in pediatric mental health and learning disorders’, Sci Data, 4: 170181.

2. Anderson, Kevin M, Tian Ge, Ru Kong, Lauren M Patrick, R Nathan Spreng, Mert R Sabuncu, BT Thomas Yeo, and Avram J Holmes. 2021. ‘Heritability of individualized cortical network topography’, Proceedings of the National Academy of Sciences, 118.

3. Arndt, Stephan, Gregg Cohen, Randall J Alliger, Victor W Swayze II, and Nancy C Andreasen. 1991. ‘Problems with ratio and proportion measures of imaged cerebral structures’, Psychiatry Research: Neuroimaging, 40: 79–89.

4. Baum, Graham L, Zaixu Cui, David R Roalf, Rastko Ciric, Richard F Betzel, Bart Larsen, Matthew Cieslak, Philip A Cook, Cedric H Xia, and Tyler M Moore. 2020. ‘Development of structure–function coupling in human brain networks during youth’, Proceedings of the National Academy of Sciences, 117: 771–78.

5. Benjamini, Yoav, and Yosef Hochberg. 1995. ‘Controlling the False Discovery Rate - a Practical and Powerful Approach to Multiple Testing’, Journal of the Royal Statistical Society Series B-Statistical Methodology, 57: 289–300.

6. Bethlehem, Richard AI, Jakob Seidlitz, Simon R White, Jacob W Vogel, Kevin M Anderson, Chris Adamson, Sophie Adler, George S Alexopoulos, Evdokia Anagnostou, and Ariosky Areces-Gonzalez. 2021. ‘Brain charts for the human lifespan’, bioRxiv.

7. Blakemore, S. J. 2012. ‘Imaging brain development: the adolescent brain’, Neuroimage, 61: 397–406.

8. Buckner, Randy L, Denise Head, Jamie Parker, Anthony F Fotenos, Daniel Marcus, John C Morris, and Abraham Z Snyder. 2004. ‘A unified approach for morphometric and functional data analysis in young, old, and demented adults using automated atlas-based head size normalization: reliability and validation against manual measurement of total intracranial volume’, Neuroimage, 23: 724–38.

9. Bzdok, Danilo, and BT Thomas Yeo. 2017. ‘Inference in the age of big data: Future perspectives on neuroscience’, Neuroimage, 155: 549–64.

10. Carlozzi, N. E., D. S. Tulsky, T. J. Wolf, S. Goodnight, R. K. Heaton, K. B. Casaletto, A. W. K. Wong, C. M. Baum, R. C. Gershon, and A. W. Heinemann. 2017. ‘Construct validity of the NIH Toolbox Cognition Battery in individuals with stroke’, Rehabil Psychol, 62: 443–54.

11. Casey, B. J., T. Cannonier, M. I. Conley, A. O. Cohen, D. M. Barch, M. M. Heitzeg, M. E. Soules, T. Teslovich, D. V. Dellarco, H. Garavan, C. A. Orr, T. D. Wager, M. T. Banich, N. K. Speer, M. T. Sutherland, M. C. Riedel, A. S. Dick, J. M. Bjork, K. M. Thomas, B. Chaarani, M. H. Mejia, D. J. Hagler, Jr., M. Daniela Cornejo, C. S. Sicat, M. P. Harms, N. U. F. Dosenbach, M. Rosenberg, E. Earl, H. Bartsch, R. Watts, J. R. Polimeni, J. M. Kuperman, D. A. Fair, A. M. Dale, and Abcd Imaging Acquisition Workgroup. 2018. ‘The Adolescent Brain Cognitive Development (ABCD) study: Imaging acquisition across 21 sites’, Dev Cogn Neurosci, 32: 43–54.

12. Casey, B. J., N. Tottenham, C. Liston, and S. Durston. 2005. ‘Imaging the developing brain: what have we learned about cognitive development?’, Trends Cogn Sci, 9: 104–10.

13. Caspi, Yaron, Rachel M Brouwer, Hugo G Schnack, Marieke E van de Nieuwenhuijzen, Wiepke Cahn, René S Kahn, Wiro J Niessen, Aad van der Lugt, and Hilleke Hulshoff Pol. 2020. ‘Changes in the intracranial volume from early adulthood to the sixth decade of life: A longitudinal study’, Neuroimage, 220: 116842.

14. Chekroud, Adam M, Emily J Ward, Monica D Rosenberg, and Avram J Holmes. 2016. ‘Patterns in the human brain mosaic discriminate males from females’, Proceedings of the National Academy of Sciences, 113: E1968–E68.

15. Chen, Jianzhong, Angela Tam, Valeria Kebets, Csaba Orban, Leon Qi Rong Ooi, Scott Marek, Nico Dosenbach, Simon Eickhoff, Danilo Bzdok, Avram J Holmes, and B T Thomas Yeo. 2020. ‘Shared and unique brain network features predict cognition, personality and mental health in childhood’, bioRxiv.

16. Cosgrove, K. P., C. M. Mazure, and J. K. Staley. 2007. ‘Evolving knowledge of sex differences in brain structure, function, and chemistry’, Biol Psychiatry, 62: 847–55.

17. Cummings, Kaitlin K, Katherine E Lawrence, Leanna M Hernandez, Emily T Wood, Susan Y Bookheimer, Mirella Dapretto, and Shulamite A Green. 2020. ‘Sex Differences in Salience Network Connectivity and its Relationship to Sensory Over-Responsivity in Youth with Autism Spectrum Disorder’, Autism Research, 13: 1489–500.

18. Dale, A. M., B. Fischl, and M. I. Sereno. 1999. ‘Cortical surface-based analysis. I. Segmentation and surface reconstruction’, Neuroimage, 9: 179–94.

19. De Bellis, M. D., M. S. Keshavan, S. R. Beers, J. Hall, K. Frustaci, A. Masalehdan, J. Noll, and A. M. Boring. 2001. ‘Sex differences in brain maturation during childhood and adolescence’, Cereb Cortex, 11: 552–7.

20. Dhamala, E., K. W. Jamison, A. Jaywant, S. Dennis, and A. Kuceyeski. 2021. ‘Distinct functional and structural connections predict crystallised and fluid cognition in healthy adults’, Hum Brain Mapp, 42: 3102–18.

21. Dhamala, E., K. W. Jamison, A. Jaywant, and A. Kuceyeski. 2021. ‘Shared functional connections within and between cortical networks predict cognitive abilities in adult males and females’, Hum Brain Mapp.

22. Dong, HaoMing, F Xavier Castellanos, Ning Yang, Zhe Zhang, Quan Zhou, Ye He, Lei Zhang, Ting Xu, Avram J Holmes, and BT Thomas Yeo. 2020. ‘Charting brain growth in tandem with brain templates at school age’, Science Bulletin, 65: 1924–34.

23. Dong, HaoMing, Daniel S Margulies, Xi-Nian Zuo, and Avram Holmes. 2020. ‘Shifting gradients of macroscale cortical organization mark the transition from childhood to adolescence’, bioRxiv.

24. Ehrlich, Stefan, Stefan Brauns, Anastasia Yendiki, Beng-Choon Ho, Vince Calhoun, S Charles Schulz, Randy L Gollub, and Scott R Sponheim. 2012. ‘Associations of cortical thickness and cognition in patients with schizophrenia and healthy controls’, Schizophrenia bulletin, 38: 1050–62.

25. Fair, D. A., A. L. Cohen, J. D. Power, N. U. Dosenbach, J. A. Church, F. M. Miezin, B. L. Schlaggar, and S. E. Petersen. 2009. ‘Functional brain networks develop from a “local to distributed” organization’, PLoS Comput Biol, 5: e1000381.

26. Farias, Sarah Tomaszewski, Dan Mungas, Bruce Reed, Owen Carmichael, Laurel Beckett, Danielle Harvey, John Olichney, Amanda Simmons, and Charles DeCarli. 2012. ‘Maximal brain size remains an important predictor of cognition in old age, independent of current brain pathology’, Neurobiology of aging, 33: 1758–68.

27. Ge, Tian, Avram J Holmes, Randy L Buckner, Jordan W Smoller, and Mert R Sabuncu. 2017. ‘Heritability analysis with repeat measurements and its application to resting-state functional connectivity’, Proceedings of the National Academy of Sciences, 114: 5521–26.

28. Gershon, R. C., M. V. Wagster, H. C. Hendrie, N. A. Fox, K. F. Cook, and C. J. Nowinski. 2013. ‘NIH toolbox for assessment of neurological and behavioral function’, Neurology, 80: S2–6.

29. Giedd, J. N., and J. L. Rapoport. 2010. ‘Structural MRI of pediatric brain development: what have we learned and where are we going?’, Neuron, 67: 728–34.

30. Gignac, Gilles E, and Timothy C Bates. 2017. ‘Brain volume and intelligence: The moderating role of intelligence measurement quality’, Intelligence, 64: 18–29.

31. Glasser, M. F., S. N. Sotiropoulos, J. A. Wilson, T. S. Coalson, B. Fischl, J. L. Andersson, J. Xu, S. Jbabdi, M. Webster, J. R. Polimeni, D. C. Van Essen, M. Jenkinson, and W. U-Minn HCP Consortium. 2013. ‘The minimal preprocessing pipelines for the Human Connectome Project’, Neuroimage, 80: 105–24.

32. Gogtay, Nitin, and Paul M Thompson. 2010. ‘Mapping gray matter development: implications for typical development and vulnerability to psychopathology’, Brain and Cognition, 72: 6–15.

33. Gong, G., Y. He, and A. C. Evans. 2011. ‘Brain connectivity: gender makes a difference’, Neuroscientist, 17: 575–91.

34. Good, Catriona D, Ingrid S Johnsrude, John Ashburner, Richard NA Henson, Karl J Friston, and Richard SJ Frackowiak. 2001. ‘A voxel-based morphometric study of ageing in 465 normal adult human brains’, Neuroimage, 14: 21–36.

35. Greene, Abigail S, Siyuan Gao, Dustin Scheinost, and R Todd Constable. 2018. ‘Task-induced brain state manipulation improves prediction of individual traits’, Nature communications, 9: 1–13.

36. Grydeland, Håkon, Petra E Vértes, František Váša, Rafael Romero-Garcia, Kirstie Whitaker, Aaron F Alexander-Bloch, Atle Bjørnerud, Ameera X Patel, Donatas Sederevič us, and Christian K Tamnes. 2019. ‘Waves of maturation and senescence in micro-structural MRI markers of human cortical myelination over the lifespan’, Cerebral cortex, 29: 1369–81.

37. Gur, R. C., and R. E. Gur. 2017. ‘Complementarity of sex differences in brain and behavior: From laterality to multimodal neuroimaging’, J Neurosci Res, 95: 189–99.

38. Gur, R. E., and R. C. Gur. 2016. ‘Sex differences in brain and behavior in adolescence: Findings from the Philadelphia Neurodevelopmental Cohort’, Neurosci Biobehav Rev, 70: 159–70.

39. Gur, Ruben C, Bruce I Turetsky, Mie Matsui, Michelle Yan, Warren Bilker, Paul Hughett, and Raquel E Gur. 1999. ‘Sex differences in brain gray and white matter in healthy young adults: correlations with cognitive performance’, Journal of Neuroscience, 19: 4065–72.

40. Hagler, D. J., Jr., S. Hatton, M. D. Cornejo, C. Makowski, D. A. Fair, A. S. Dick, M. T. Sutherland, B. J. Casey, D. M. Barch, M. P. Harms, R. Watts, J. M. Bjork, H. P. Garavan, L. Hilmer, C. J. Pung, C. S. Sicat, J. Kuperman, H. Bartsch, F. Xue, M. M. Heitzeg, A. R. Laird, T. T. Trinh, R. Gonzalez, S. F. Tapert, M. C. Riedel, L. M. Squeglia, L. W. Hyde, M. D. Rosenberg, E. A. Earl, K. D. Howlett, F. C. Baker, M. Soules, J. Diaz, O. R. de Leon, W. K. Thompson, M. C. Neale, M. Herting, E. R. Sowell, R. P. Alvarez, S. W. Hawes, M. Sanchez, J. Bodurka, F. J. Breslin, A. S. Morris, M. P. Paulus, W. K. Simmons, J. R. Polimeni, A. van der Kouwe, A. S. Nencka, K. M. Gray, C. Pierpaoli, J. A. Matochik, A. Noronha, W. M. Aklin, K. Conway, M. Glantz, E. Hoffman, R. Little, M. Lopez, V. Pariyadath, S. R. Weiss, D. L. Wolff-Hughes, R. DelCarmen-Wiggins, S. W. Feldstein Ewing, O. Miranda-Dominguez, B. J. Nagel, A. J. Perrone, D. T. Sturgeon, A. Goldstone, A. Pfefferbaum, K. M. Pohl, D. Prouty, K. Uban, S. Y. Bookheimer, M. Dapretto, A. Galvan, K. Bagot, J. Giedd, M. A. Infante, J. Jacobus, K. Patrick, P. D. Shilling, R. Desikan, Y. Li, L. Sugrue, M. T. Banich, N. Friedman, J. K. Hewitt, C. Hopfer, J. Sakai, J. Tanabe, L. B. Cottler, S. J. Nixon, L. Chang, C. Cloak, T. Ernst, G. Reeves, D. N. Kennedy, S. Heeringa, S. Peltier, J. Schulenberg, C. Sripada, R. A. Zucker, W. G. Iacono, M. Luciana, F. J. Calabro, D. B. Clark, D. A. Lewis, B. Luna, C. Schirda, T. Brima, J. J. Foxe, E. G. Freedman, D. W. Mruzek, M. J. Mason, R. Huber, E. McGlade, A. Prescot, P. F. Renshaw, D. A. Yurgelun-Todd, N. A. Allgaier, J. A. Dumas, M. Ivanova, A. Potter, P. Florsheim, C. Larson, K. Lisdahl, M. E. Charness, B. Fuemmeler, J. M. Hettema, H. H. Maes, J. Steinberg, A. P. Anokhin, P. Glaser, A. C. Heath, P. A. Madden, A. Baskin-Sommers, R. T. Constable, S. J. Grant, G. J. Dowling, S. A. Brown, T. L. Jernigan, and A. M. Dale. 2019. ‘Image processing and analysis methods for the Adolescent Brain Cognitive Development Study’, Neuroimage, 202: 116091.

41. Hagmann, P., O. Sporns, N. Madan, L. Cammoun, R. Pienaar, V. J. Wedeen, R. Meuli, J. P. Thiran, and P. E. Grant. 2010. ‘White matter maturation reshapes structural connectivity in the late developing human brain’, Proc Natl Acad Sci U S A, 107: 19067–72.

42. Hartberg, CB, K Sundet, LM Rimol, UK Haukvik, EH Lange, R Nesvåg, AM Dale, I Melle, OA Andreassen, and I Agartz. 2011. ‘Brain cortical thickness and surface area correlates of neurocognitive performance in patients with schizophrenia, bipolar disorder, and healthy adults’, Journal of the International Neuropsychological Society, 17: 1080–93.

43. Haufe, Stefan, Frank Meinecke, Kai Görgen, Sven Dähne, John-Dylan Haynes, Benjamin Blankertz, and Felix Bießmann. 2014. ‘On the interpretation of weight vectors of linear models in multivariate neuroimaging’, Neuroimage, 87: 96–110.

44. He, Tong, Lijun An, Jiashi Feng, Danilo Bzdok, Avram J Holmes, Simon B Eickhoff, and Boon Thye Thomas Yeo. 2020. ‘Meta-matching: a simple framework to translate phenotypic predictive models from big to small data’, bioRxiv.

45. He, Tong, Ru Kong, Avram J Holmes, Minh Nguyen, Mert R Sabuncu, Simon B Eickhoff, Danilo Bzdok, Jiashi Feng, and BT Thomas Yeo. 2020. ‘Deep neural networks and kernel regression achieve comparable accuracies for functional connectivity prediction of behavior and demographics’, Neuroimage, 206: 116276.

46. Heaton, R. K., N. Akshoomoff, D. Tulsky, D. Mungas, S. Weintraub, S. Dikmen, J. Beaumont, K. B. Casaletto, K. Conway, J. Slotkin, and R. Gershon. 2014. ‘Reliability and validity of composite scores from the NIH Toolbox Cognition Battery in adults’, J Int Neuropsychol Soc, 20: 588–98.

47. Hill, Jason, Terrie Inder, Jeffrey Neil, Donna Dierker, John Harwell, and David Van Essen. 2010. ‘Similar patterns of cortical expansion during human development and evolution’, Proceedings of the National Academy of Sciences, 107: 13135–40.

48. Holmes, Avram J, Marisa O Hollinshead, Timothy M O’keefe, Victor I Petrov, Gabriele R Fariello, Lawrence L Wald, Bruce Fischl, Bruce R Rosen, Ross W Mair, and Joshua L Roffman. 2015. ‘Brain Genomics Superstruct Project initial data release with structural, functional, and behavioral measures’, Scientific data, 2: 1–16.

49. Ingalhalikar, M., A. Smith, D. Parker, T. D. Satterthwaite, M. A. Elliott, K. Ruparel, H. Hakonarson, R. E. Gur, R. C. Gur, and R. Verma. 2014. ‘Sex differences in the structural connectome of the human brain’, Proc Natl Acad Sci U S A, 111: 823–8.

50. Jäncke, Lutz. 2018. ‘Sex/gender differences in cognition, neurophysiology, and neuroanatomy’, F1000Research, 7.

51. Jiang, Rongtao, Vince D Calhoun, Yue Cui, Shile Qi, Chuanjun Zhuo, Jin Li, Rex Jung, Jian Yang, Yuhui Du, and Tianzi Jiang. 2020. ‘Multimodal data revealed different neurobiological correlates of intelligence between males and females’, Brain imaging and behavior, 14: 1979–93.

52. Jiang, Rongtao, Vince D Calhoun, Lingzhong Fan, Nianming Zuo, Rex Jung, Shile Qi, Dongdong Lin, Jin Li, Chuanjun Zhuo, Ming Song, Zening Fu, Tianzi Jiang, and Jing Sui. 2020. ‘Gender differences in connectome-based predictions of individualized intelligence quotient and sub-domain scores’, Cerebral cortex, 30: 888–900.

53. Joel, Daphna, Zohar Berman, Ido Tavor, Nadav Wexler, Olga Gaber, Yaniv Stein, Nisan Shefi, Jared Pool, Sebastian Urchs, and Daniel S Margulies. 2015. ‘Sex beyond the genitalia: The human brain mosaic’, Proceedings of the National Academy of Sciences, 112: 15468–73.

54. Kohoutová, Lada, Juyeon Heo, Sungmin Cha, Sungwoo Lee, Taesup Moon, Tor D Wager, and Choong-Wan Woo. 2020. ‘Toward a unified framework for interpreting machine-learning models in neuroimaging’, Nature protocols, 15: 1399–435.

55. Kong, Ru, Qing Yang, Evan Gordon, Aihuiping Xue, Xiaoxuan Yan, Csaba Orban, Xi-Nian Zuo, Nathan Spreng, Tian Ge, Avram Holmes, Simon Eickhoff, and B T Thomas Yeo. 2021. ‘Individual-Specific Areal-Level Parcellations Improve Functional Connectivity Prediction of Behavior’, Cerebral cortex.

56. Krogsrud, Stine K, Athanasia M Mowinckel, Donatas Sederevicius, Didac Vidal-Piñeiro, Inge K Amlien, Yunpeng Wang, Øystein Sørensen, Kristine B Walhovd, and Anders M Fjell. 2021. ‘Relationships between apparent cortical thickness and working memory across the lifespan-Effects of genetics and socioeconomic status’, Developmental cognitive neuroscience, 51: 100997.

57. Lenroot, R. K., and J. N. Giedd. 2010. ‘Sex differences in the adolescent brain’, Brain Cogn, 72: 46–55.

58. Li, J. W., R. Kong, R. Liegeois, C. Orban, Y. R. Tan, N. B. Sun, A. J. Holmes, M. R. Sabuncu, T. Ge, and B. T. T. Yeo. 2019. ‘Global signal regression strengthens association between resting-state functional connectivity and behavior’, Neuroimage, 196: 126–41.

59. Liu, Dawei, Hans J Johnson, Jeffrey D Long, Vincent A Magnotta, and Jane S Paulsen. 2014. ‘The power-proportion method for intracranial volume correction in volumetric imaging analysis’, Frontiers in Neuroscience, 8: 356.

60. MacKinnon, James G. 2009. ‘Bootstrap hypothesis testing’, Handbook of computational econometrics, 183: 213.

61. MacLullich, AMJ, KJ Ferguson, IJ Deary, JR Seckl, JM Starr, and JM Wardlaw. 2002. ‘Intracranial capacity and brain volumes are associated with cognition in healthy elderly men’, Neurology, 59: 169–74.

62. Madan, Christopher R. 2021. ‘Scan once, analyse many: using large open-access neuroimaging datasets to understand the brain’, Neuroinformatics: 1–29.

63. Mansour, Sina L, Ye Tian, BT Thomas Yeo, Vanessa Cropley, and Andrew Zalesky. 2021. ‘High-resolution connectomic fingerprints: Mapping neural identity and behavior’, Neuroimage, 229: 117695.

64. Marcus, D. S., M. P. Harms, A. Z. Snyder, M. Jenkinson, J. A. Wilson, M. F. Glasser, D. M. Barch, K. A. Archie, G. C. Burgess, M. Ramaratnam, M. Hodge, W. Horton, R. Herrick, T. Olsen, M. McKay, M. House, M. Hileman, E. Reid, J. Harwell, T. Coalson, J. Schindler, J. S. Elam, S. W. Curtiss, D. C. Van Essen, and W. U-Minn HCP Consortium. 2013. ‘Human Connectome Project informatics: quality control, database services, and data visualization’, Neuroimage, 80: 202–19.

65. Marek, Scott, Brenden Tervo-Clemmens, Finnegan J Calabro, David F Montez, Benjamin P Kay, Alexander S Hatoum, Meghan Rose Donohue, William Foran, Ryland L Miller, and Eric Feczko. 2020. ‘Towards reproducible brain-wide association studies’, bioRxiv.

66. Mathalon, Daniel H, Edith V Sullivan, Jody M Rawles, and Adolf Pfefferbaum. 1993. ‘Correction for head size in brain-imaging measurements’, Psychiatry Research: Neuroimaging, 50: 121–39.

67. Matsumae, Mitsunori, Ron Kikinis, István A Mórocz, Antonio V Lorenzo, Tamás Sándor, Marilyn S Albert, Peter McL Black, and Ferenc A Jolesz. 1996. ‘Age-related changes in intracranial compartment volumes in normal adults assessed by magnetic resonance imaging’, Journal of neurosurgery, 84: 982–91.

68. Mills, Kathryn L, Anne-Lise Goddings, Megan M Herting, Rosa Meuwese, Sarah-Jayne Blakemore, Eveline A Crone, Ronald E Dahl, Berna Güroğlu, Armin Raznahan, and Elizabeth R Sowell. 2016. ‘Structural brain development between childhood and adulthood: Convergence across four longitudinal samples’, Neuroimage, 141: 273–81.

69. Mungas, D., R. Heaton, D. Tulsky, P. D. Zelazo, J. Slotkin, D. Blitz, J. S. Lai, and R. Gershon. 2014. ‘Factor structure, convergent validity, and discriminant validity of the NIH Toolbox Cognitive Health Battery (NIHTB-CHB) in adults’, J Int Neuropsychol Soc, 20: 579–87.

70. Ooi, Leon Qi Rong, Jianzhong Chen, Shaoshi Zhang, Ru Kong, Jingwei Li, Elvisha Dhamala, Juan Helen Zhou, Avram Holmes, and B. T. Thomas Yeo. 2022. ‘Comparison of individualized behavioral predictions across anatomical, diffusion and functional connectivity MRI’, bioRxiv.

71. Parkes, Linden, Tyler M Moore, Monica E Calkins, Philip A Cook, Matthew Cieslak, David R Roalf, Daniel H Wolf, Ruben C Gur, Raquel E Gur, and Theodore D Satterthwaite. 2021. ‘Transdiagnostic dimensions of psychopathology explain individuals’ unique deviations from normative neurodevelopment in brain structure’, Translational psychiatry, 11: 1–13.

72. Pintzka, Carl Wolfgang S, Tor Ivar Hansen, Hallvard R Evensmoen, and Asta Kristine Håberg. 2015. ‘Marked effects of intracranial volume correction methods on sex differences in neuroanatomical structures: a HUNT MRI study’, Frontiers in Neuroscience, 9: 238.

73. Rosenberg, Monica D, BJ Casey, and Avram J Holmes. 2018. ‘Prediction complements explanation in understanding the developing brain’, Nature communications, 9: 1–13.

74. Sanchis-Segura, Carla, Maria Victoria Ibañez-Gual, Jesús Adrián-Ventura, Naiara Aguirre, Álvaro Javier Gómez-Cruz, César Avila, and Cristina Forn. 2019. ‘Sex differences in gray matter volume: how many and how large are they really?’, Biology of sex Differences, 10: 1–19.

75. Sanchis-Segura, Carla, Maria Victoria Ibañez-Gual, Naiara Aguirre, Álvaro Javier Cruz-Gómez, and Cristina Forn. 2020. ‘Effects of different intracranial volume correction methods on univariate sex differences in grey matter volume and multivariate sex prediction’, Scientific reports, 10: 1–15.

76. Sanfilipo, Michael P, Ralph HB Benedict, Robert Zivadinov, and Rohit Bakshi. 2004. ‘Correction for intracranial volume in analysis of whole brain atrophy in multiple sclerosis: the proportion vs. residual method’, Neuroimage, 22: 1732–43.

77. Satterthwaite, T. D., M. A. Elliott, K. Ruparel, J. Loughead, K. Prabhakaran, M. E. Calkins, R. Hopson, C. Jackson, J. Keefe, M. Riley, F. D. Mentch, P. Sleiman, R. Verma, C. Davatzikos, H. Hakonarson, R. C. Gur, and R. E. Gur. 2014. ‘Neuroimaging of the Philadelphia neurodevelopmental cohort’, Neuroimage, 86: 544–53.

78. Satterthwaite, T. D., D. H. Wolf, D. R. Roalf, K. Ruparel, G. Erus, S. Vandekar, E. D. Gennatas, M. A. Elliott, A. Smith, H. Hakonarson, R. Verma, C. Davatzikos, R. E. Gur, and R. C. Gur. 2015. ‘Linked Sex Differences in Cognition and Functional Connectivity in Youth’, Cereb Cortex, 25: 2383–94.

79. Schaefer, Alexander, Ru Kong, Evan M Gordon, Timothy O Laumann, Xi-Nian Zuo, Avram J Holmes, Simon B Eickhoff, and BT Thomas Yeo. 2018. ‘Local-global parcellation of the human cerebral cortex from intrinsic functional connectivity MRI’, Cerebral cortex, 28: 3095–114.

80. Scheinost, Dustin, Emily S Finn, Fuyuze Tokoglu, Xilin Shen, Xenophon Papademetris, Michelle Hampson, and R Todd Constable. 2015. ‘Sex differences in normal age trajectories of functional brain networks’, Human Brain Mapping, 36: 1524–35.

81. Scheinost, Dustin, Stephanie Noble, Corey Horien, Abigail S Greene, Evelyn MR Lake, Mehraveh Salehi, Siyuan Gao, Xilin Shen, David O’Connor, and Daniel S Barron. 2019. ‘Ten simple rules for predictive modeling of individual differences in neuroimaging’, Neuroimage, 193: 35–45.

82. Seidlitz, J., F. Vasa, M. Shinn, R. Romero-Garcia, K. J. Whitaker, P. E. Vertes, K. Wagstyl, P. Kirkpatrick Reardon, L. Clasen, S. Liu, A. Messinger, D. A. Leopold, P. Fonagy, R. J. Dolan, P. B. Jones, I. M. Goodyer, Nspn Consortium, A. Raznahan, and E. T. Bullmore. 2018. ‘Morphometric Similarity Networks Detect Microscale Cortical Organization and Predict Inter-Individual Cognitive Variation’, Neuron, 97: 231–47 e7.

83. Somerville, L. H., S. Y. Bookheimer, R. L. Buckner, G. C. Burgess, S. W. Curtiss, M. Dapretto, J. S. Elam, M. S. Gaffrey, M. P. Harms, C. Hodge, S. Kandala, E. K. Kastman, T. E. Nichols, B. L. Schlaggar, S. M. Smith, K. M. Thomas, E. Yacoub, D. C. Van Essen, and D. M. Barch. 2018. ‘The Lifespan Human Connectome Project in Development: A large-scale study of brain connectivity development in 5-21 year olds’, Neuroimage, 183: 456–68.

84. Stiles, J., and T. L. Jernigan. 2010. ‘The basics of brain development’, Neuropsychol Rev, 20: 327–48.

85. Sudlow, Cathie, John Gallacher, Naomi Allen, Valerie Beral, Paul Burton, John Danesh, Paul Downey, Paul Elliott, Jane Green, and Martin Landray. 2015. ‘UK biobank: an open access resource for identifying the causes of a wide range of complex diseases of middle and old age’, PLoS medicine, 12.

86. Sydnor, Valerie J, Bart Larsen, Danielle S Bassett, Aaron Alexander-Bloch, Damien A Fair, Conor Liston, Allyson P Mackey, Michael P Milham, Adam Pines, and David R Roalf. 2021. ‘Neurodevelopment of the association cortices: Patterns, mechanisms, and implications for psychopathology’, Neuron, 109: 2820–46.

87. Tian, Ye, and Andrew Zalesky. 2021. ‘Machine learning prediction of cognition from functional connectivity: Are feature weights reliable?’, bioRxiv.

88. Van Essen, D. C., S. M. Smith, D. M. Barch, T. E. Behrens, E. Yacoub, K. Ugurbil, and W. U-Minn HCP Consortium. 2013. ‘The WU-Minn Human Connectome Project: an overview’, Neuroimage, 80: 62–79.

89. Van Loenhoud, Anna Catharina, Colin Groot, Jacob William Vogel, Wiesje Maria Van Der Flier, and Rik Ossenkoppele. 2018. ‘Is intracranial volume a suitable proxy for brain reserve?’, Alzheimer’s research & therapy, 10: 1–12.

90. Voevodskaya, Olga, Andrew Simmons, Richard Nordenskjöld, Joel Kullberg, Håkan Ahlström, Lars Lind, Lars-Olof Wahlund, Elna-Marie Larsson, Eric Westman, and Alzheimer’s Disease Neuroimaging Initiative. 2014. ‘The effects of intracranial volume adjustment approaches on multiple regional MRI volumes in healthy aging and Alzheimer’s disease’, Frontiers in aging neuroscience, 6: 264.

91. Wang, Fan, Chunfeng Lian, Zhengwang Wu, Han Zhang, Tengfei Li, Yu Meng, Li Wang, Weili Lin, Dinggang Shen, and Gang Li. 2019. ‘Developmental topography of cortical thickness during infancy’, Proceedings of the National Academy of Sciences, 116: 15855–60.

92. Weintraub, S., P. J. Bauer, P. D. Zelazo, K. Wallner-Allen, S. S. Dikmen, R. K. Heaton, D. S. Tulsky, J. Slotkin, D. L. Blitz, N. E. Carlozzi, R. J. Havlik, J. L. Beaumont, D. Mungas, J. J. Manly, B. G. Borosh, C. J. Nowinski, and R. C. Gershon. 2013. ‘I. NIH Toolbox Cognition Battery (CB): introduction and pediatric data’, Monogr Soc Res Child Dev, 78: 1–15.

93. Weintraub, S., S. S. Dikmen, R. K. Heaton, D. S. Tulsky, P. D. Zelazo, J. Slotkin, N. E. Carlozzi, P. J. Bauer, K. Wallner-Allen, N. Fox, R. Havlik, J. L. Beaumont, D. Mungas, J. J. Manly, C. Moy, K. Conway, E. Edwards, C. J. Nowinski, and R. Gershon. 2014. ‘The cognition battery of the NIH toolbox for assessment of neurological and behavioral function: validation in an adult sample’, J Int Neuropsychol Soc, 20: 567–78.

94. Wierenga, L. M., M. G. N. Bos, F. van Rossenberg, and E. A. Crone. 2019. ‘Sex Effects on Development of Brain Structure and Executive Functions: Greater Variance than Mean Effects’, J Cogn Neurosci: 1–23.

95. Wierenga, Lara M, Marieke Langen, Bob Oranje, and Sarah Durston. 2014. ‘Unique developmental trajectories of cortical thickness and surface area’, Neuroimage, 87: 120–26.

96. Yarkoni, Tal, and Jacob Westfall. 2017. ‘Choosing prediction over explanation in psychology: Lessons from machine learning’, Perspectives on Psychological Science, 12: 1100–22.

97. Yeo, BT Thomas, Fenna M Krienen, Jorge Sepulcre, Mert R Sabuncu, Danial Lashkari, Marisa Hollinshead, Joshua L Roffman, Jordan W Smoller, Lilla Zöllei, and Jonathan R Polimeni. 2011. ‘The organization of the human cerebral cortex estimated by intrinsic functional connectivity’, Journal of neurophysiology, 106: 1125–65.

98. Zelazo, P. D., J. E. Anderson, J. Richler, K. Wallner-Allen, J. L. Beaumont, K. P. Conway, R. Gershon, and S. Weintraub. 2014. ‘NIH Toolbox Cognition Battery (CB): validation of executive function measures in adults’, J Int Neuropsychol Soc, 20: 620–9.

99. Zelazo, P. D., and P. J. Bauer. 2013. National Institutes of Health Toolbox cognition battery (NIH Toolbox CB): Validation for children between 3 and 15 years (Wiley Hoboken, NJ).

100. Zhou, Dongming, Catherine Lebel, Sarah Treit, Alan Evans, and Christian Beaulieu. 2015. ‘Accelerated longitudinal cortical thinning in adolescence’, Neuroimage, 104: 138–45.

