## Supplementary Materials for "Proportional intracranial volume correction differentially biases behavioral predictions across neuroanatomical features and populations"

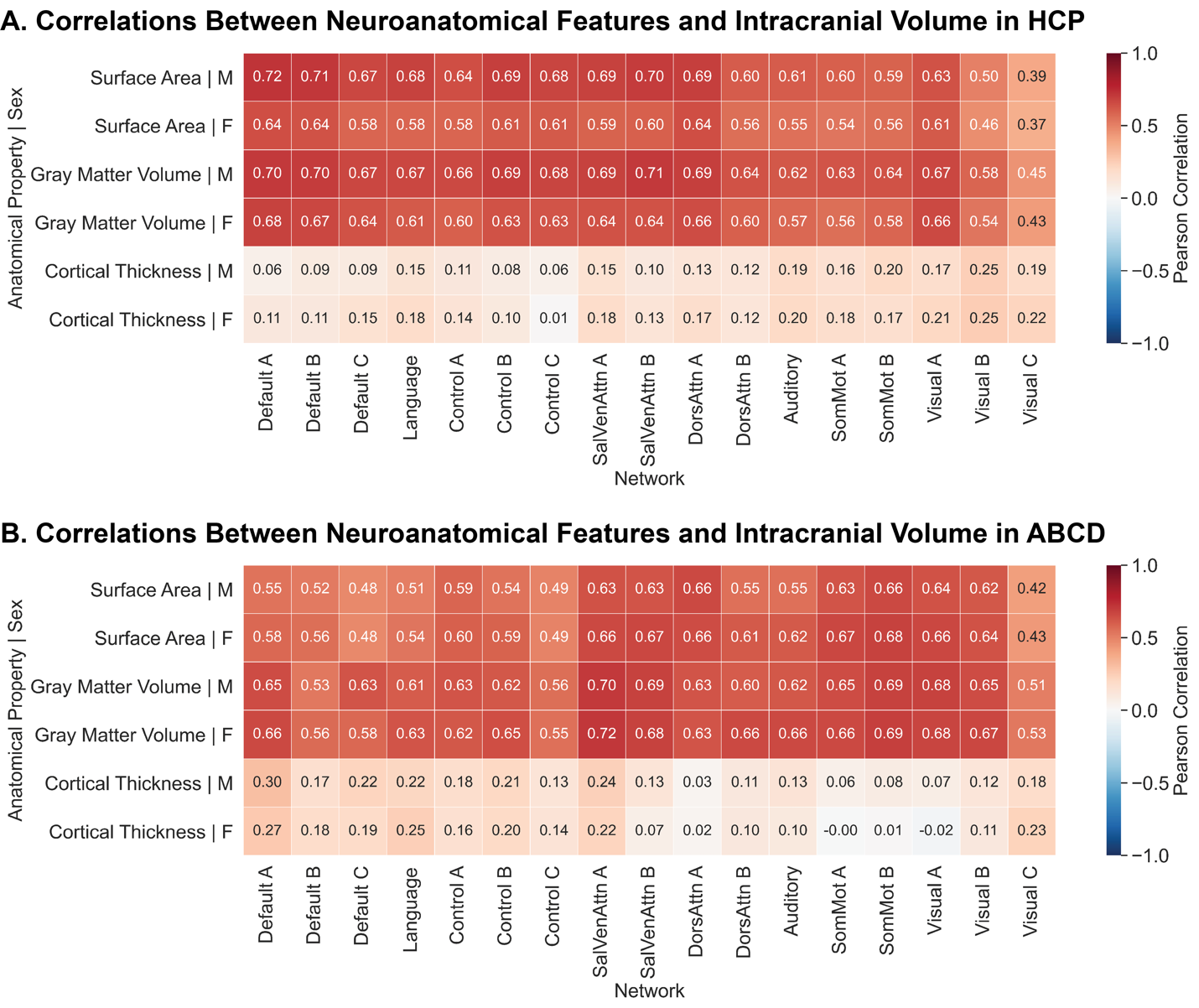

Figure S1: **Total surface area and gray matter volume, and average cortical thickness across distinct cortical networks exhibit correlations with intracranial volume.** Sex-specific correlation (Pearson’s correlation coefficient) between network-level neuroanatomical properties (total surface area, total gray matter volume, and average cortical thickness) and total intracranial volume in HCP (A) and ABCD (B). M denotes male and F denotes female. SalVenAttn – Salience/Ventral Attention; DorsAttn – Dorsal Attention; SomMot – Somatomotor. Networks are ordered from heteromodal (left) to unimodal (right).

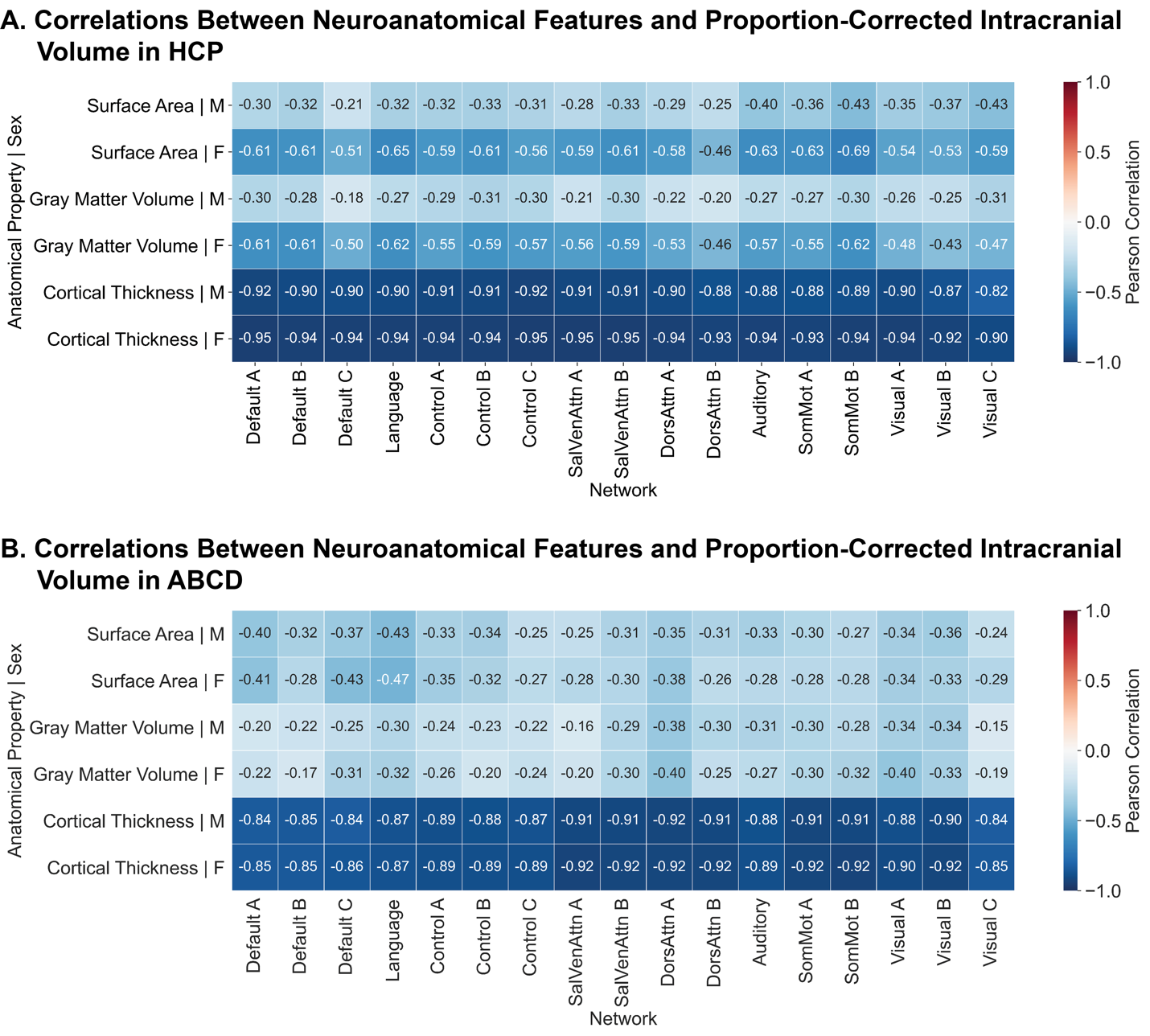

Figure S2: **Total surface area and gray matter volume, and average cortical thickness across distinct cortical networks exhibit negative correlations with proportion-corrected intracranial volume.** Sex-specific correlation (Pearson’s correlation coefficient) between network-level neuroanatomical properties (total surface area, total gray matter volume, and average cortical thickness) and total intracranial volume in HCP (A) and ABCD (B). M denotes male and F denotes female. SalVenAttn – Salience/Ventral Attention; DorsAttn – Dorsal Attention; SomMot – Somatomotor. Networks are ordered from heteromodal (left) to unimodal (right).

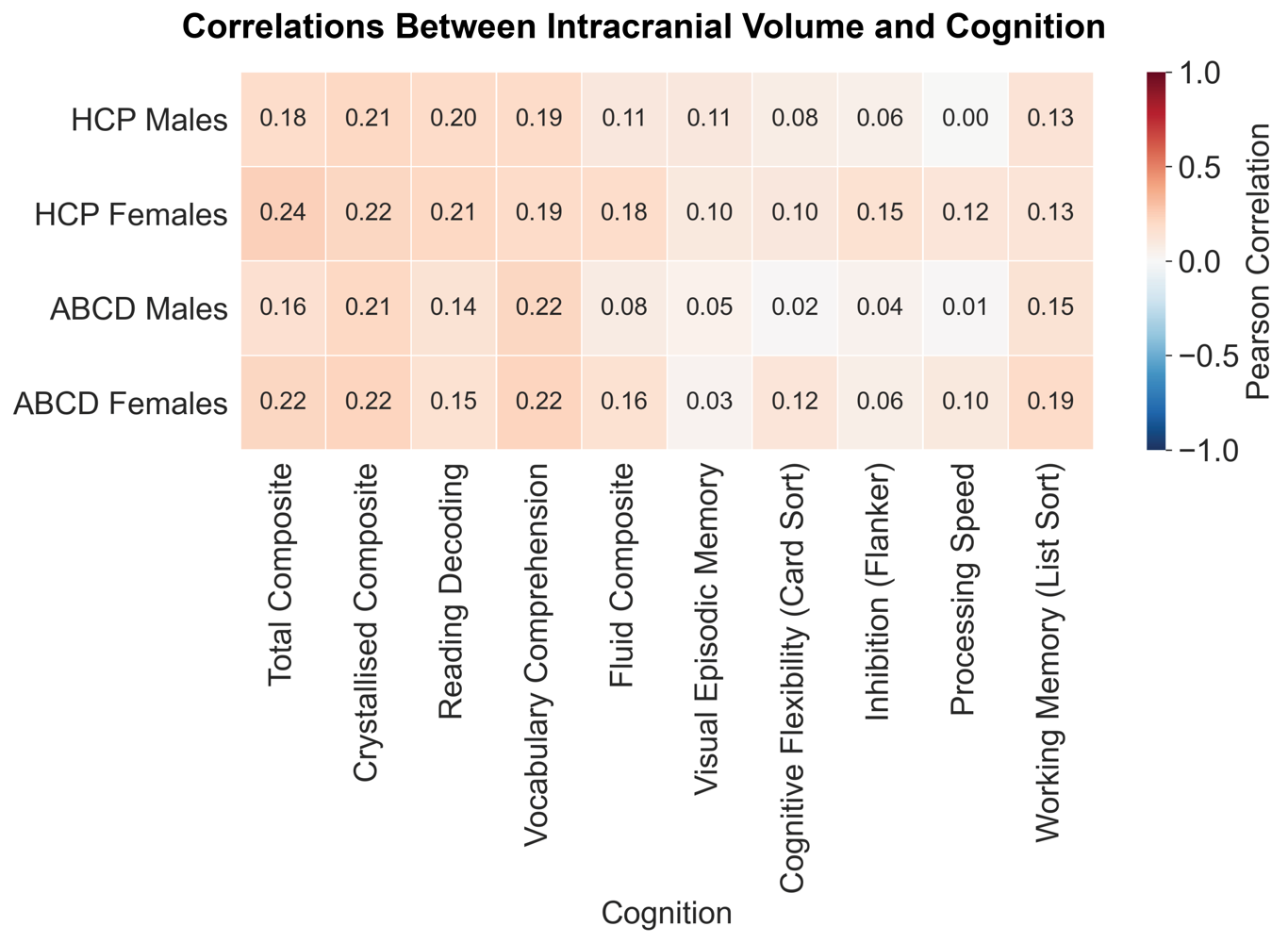

Figure S3: **Intracranial volume exhibit varying correlations with cognition.** Sex-specific correlation (Pearson’s correlation coefficient) between intracranial volume and cognitive composite and individual task scores in HCP males, HCP females, ABCD males, and ABCD females.

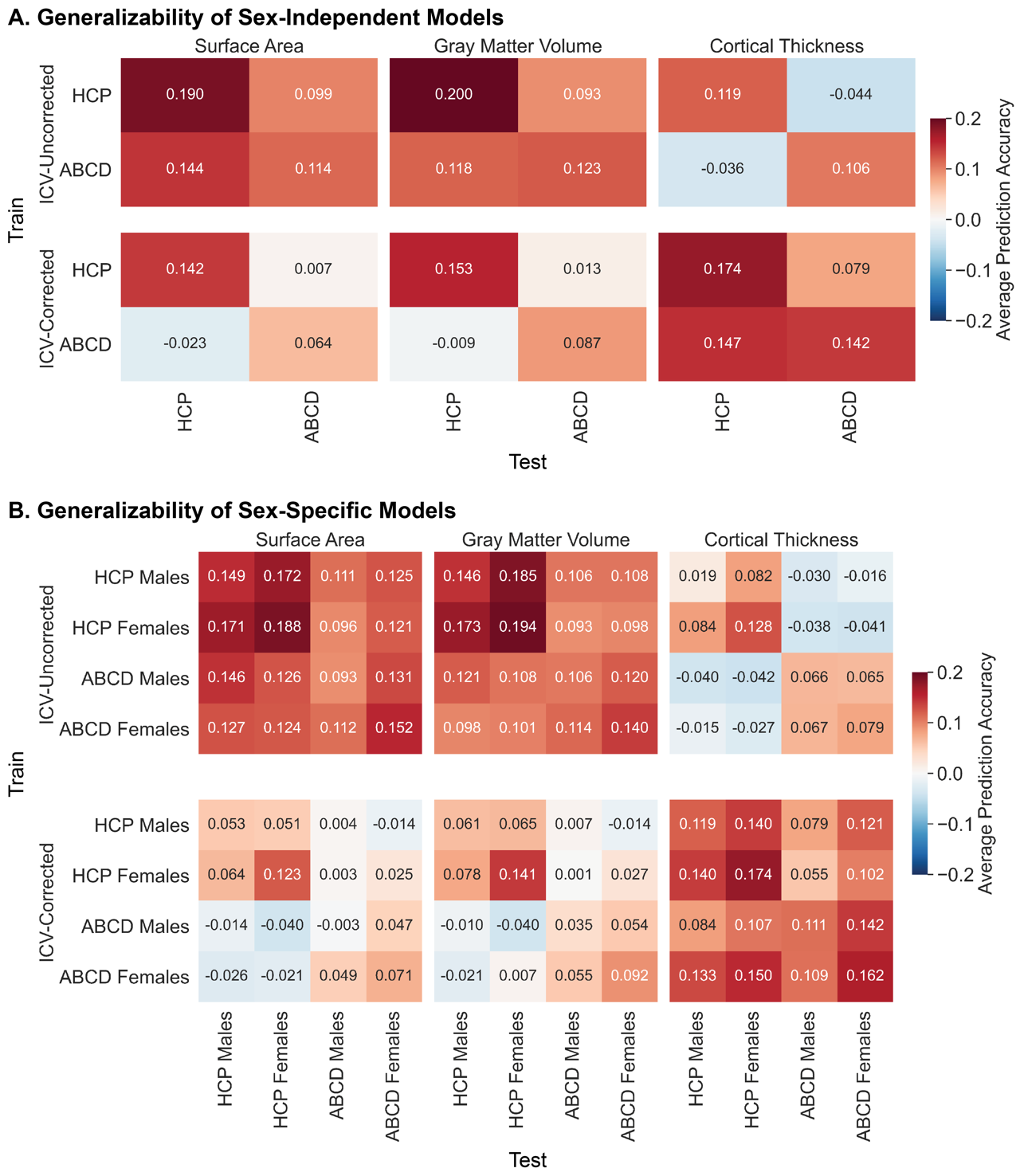

Figure S4: **Intracranial volume correction reduces generalizability of models based on surface area and gray matter volume but increases generalizability of models based on cortical thickness.** Generalizability of sex-independent (A) and sex-specific (B) models across sexes (males and females) and datasets (HCP and ABCD). Average prediction accuracies across all 10 cognitive scores based on surface area (left), gray matter volume (middle), and cortical thickness (right) using raw anatomical properties are shown in the top panels, and predictions using ICV proportion-corrected anatomical properties are shown in the bottom panels. The populations that the models were trained on are shown along the rows, and the populations that the models were tested on are shown along the columns.

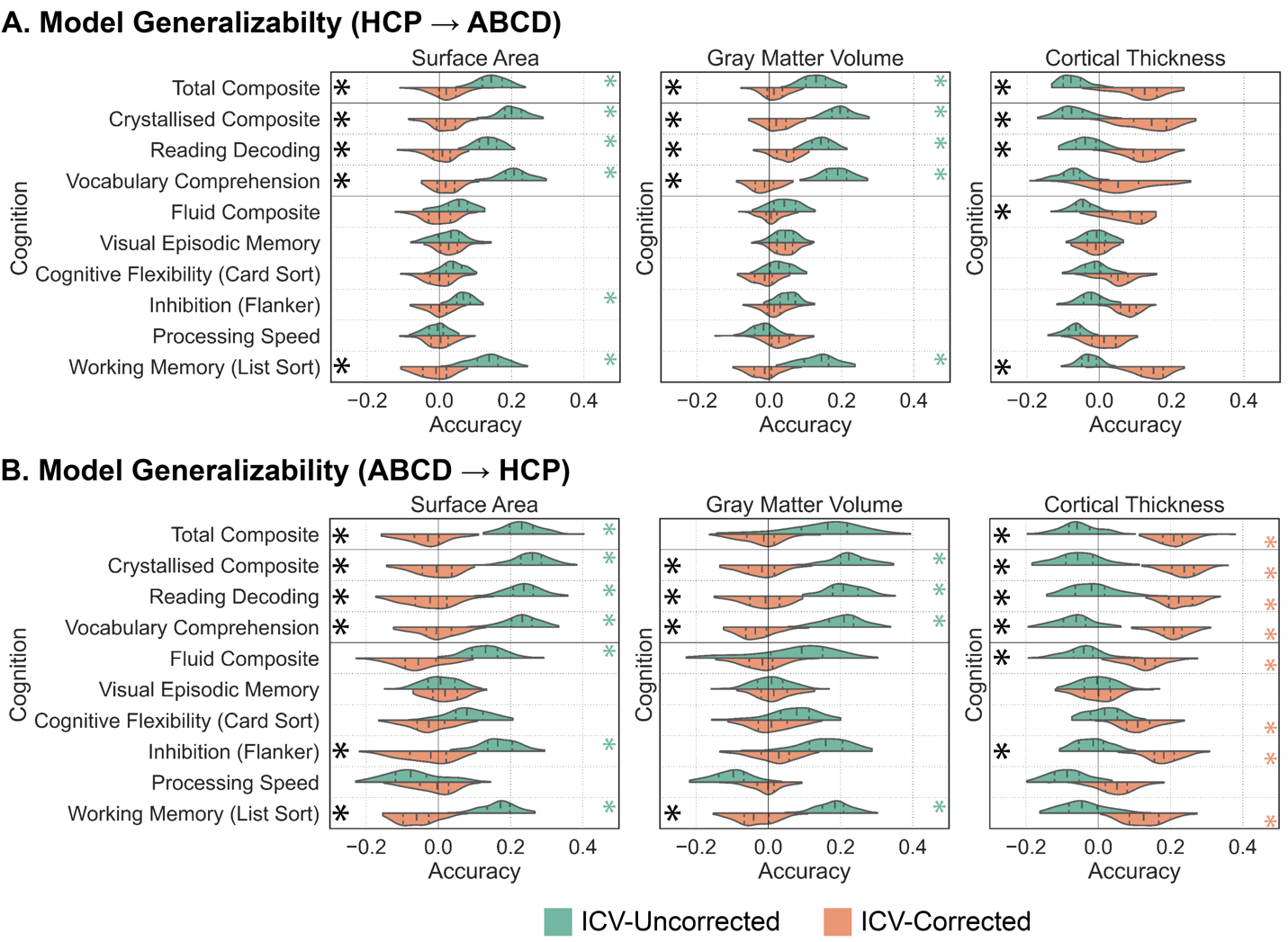

Figure S5: **Accounting for intracranial volume** **reduces model generalizability in terms of predictive accuracies of cognition based on surface area and gray matter volume, and increases predictive accuracies based on cortical thickness.** Prediction accuracies (Pearson’s correlation coefficient between observed and predicted scores) for sex-independent models predicting cognitive scores in model trained on HCP and evaluated on ABCD (A) and models trained on ABCD and evaluated on HCP (B). Predictions based on surface area (left), gray matter volume (middle), and cortical thickness (right) using raw (green) and ICV proportion-corrected (orange) anatomical properties are shown. Green and orange asterisks (*) denote that the model performed above chance levels based on permutation tests (corrected p<0.05). Black asterisks (*) denote that model performance was significantly different between the raw and ICV proportion-corrected predictions based on exact tests for differences (corrected p<0.05). The shape of the violin plots indicates the entire distribution of values, dashed lines indicate the median, and dotted lines indicate the interquartile range.

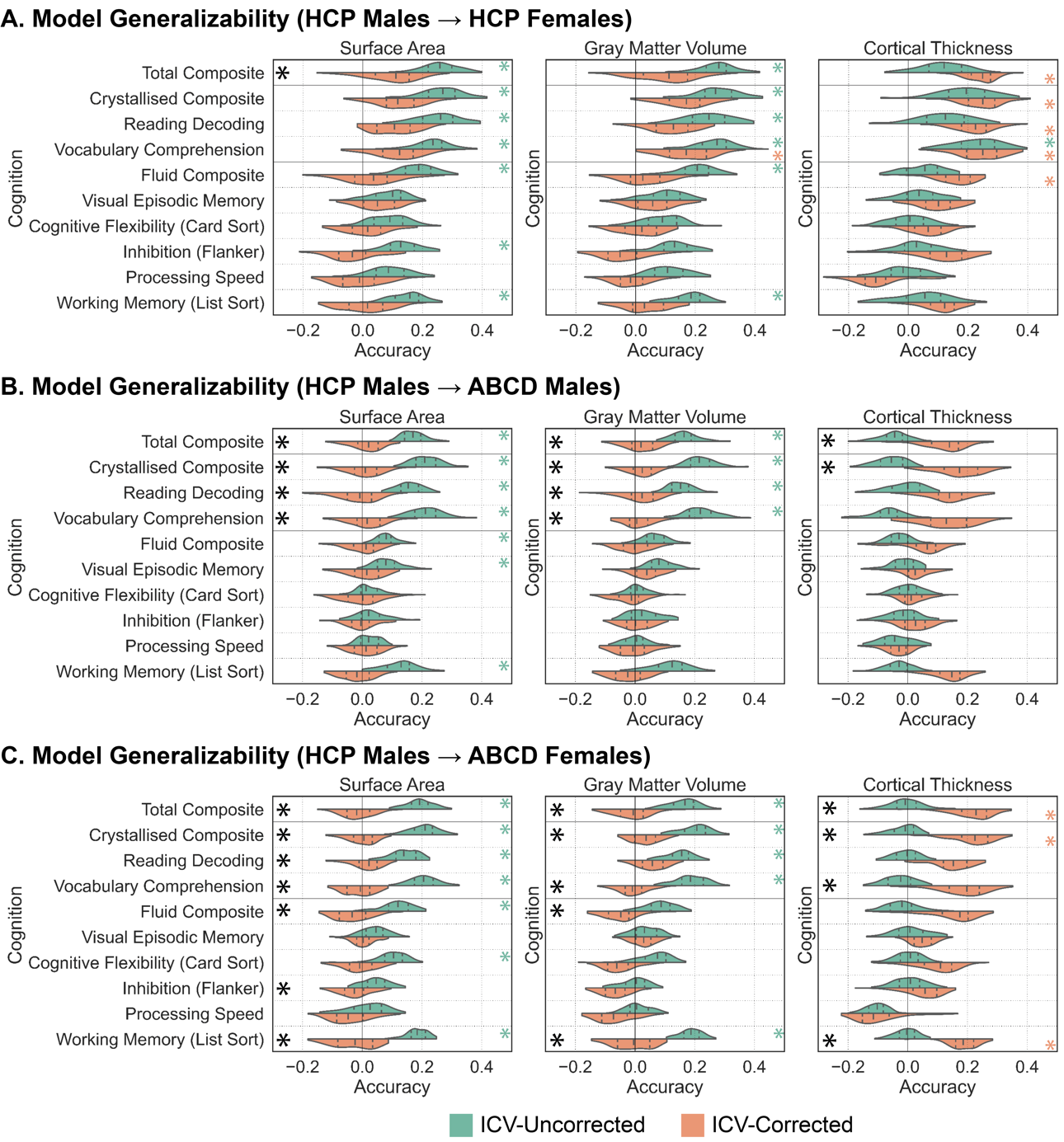

Figure S6: **Accounting for intracranial volume differentially impacts model generalizability in terms of predictive accuracies across surface area, gray matter volume, and cortical thickness in a sex specific manner.** Prediction accuracies (Pearson’s correlation coefficient between observed and predicted scores) for sex-specific models trained on HCP males to predict cognitive scores and evaluated on HCP females (A), ABCD males (B), and ABCD females (C). Predictions based on surface area (left), gray matter volume (middle), and cortical thickness (right) using raw (green) and ICV proportion-corrected (orange) anatomical properties are shown. Green and orange asterisks (*) denote that the model performed above chance levels based on permutation tests (corrected p<0.05). Black asterisks (*) denote that model performance was significantly different between the raw and ICV proportion-corrected predictions based on exact tests for differences (corrected p<0.05). The shape of the violin plots indicates the entire distribution of values, dashed lines indicate the median, and dotted lines indicate the interquartile range.

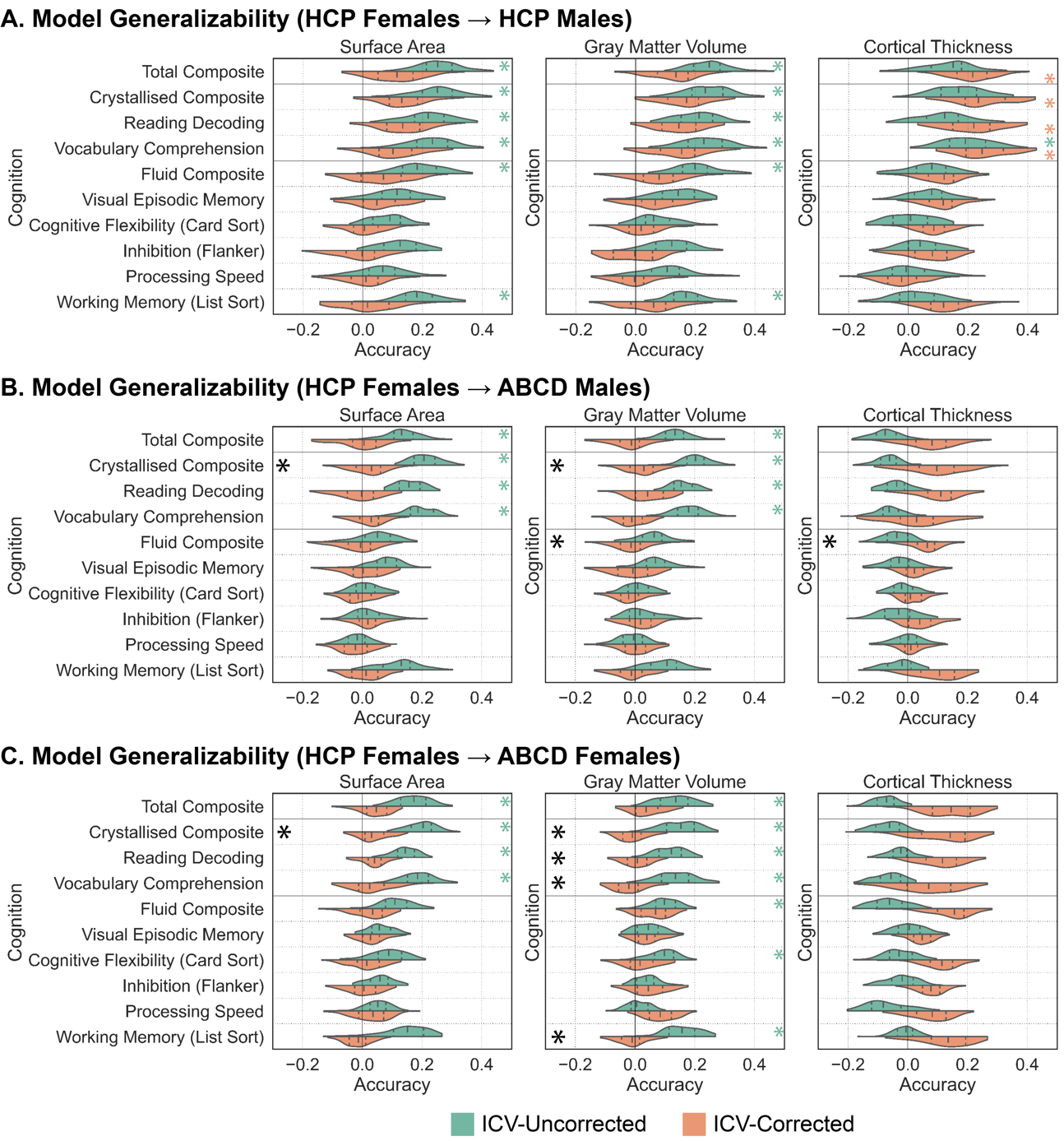

Figure S7: **Accounting for intracranial volume differentially impacts model generalizability in terms of predictive accuracies across surface area, gray matter volume, and cortical thickness in a sex specific manner.** Prediction accuracies (Pearson’s correlation coefficient between observed and predicted scores) for sex-specific models trained on HCP females to predict cognitive scores and evaluated on HCP males (A), ABCD males (B), and ABCD females (C). Predictions based on surface area (left), gray matter volume (middle), and cortical thickness (right) using raw (green) and ICV proportion-corrected (orange) anatomical properties are shown. Green and orange asterisks (*) denote that the model performed above chance levels based on permutation tests (corrected p<0.05). Black asterisks (*) denote that model performance was significantly different between the raw and ICV proportion-corrected predictions based on exact tests for differences (corrected p<0.05). The shape of the violin plots indicates the entire distribution of values, dashed lines indicate the median, and dotted lines indicate the interquartile range.

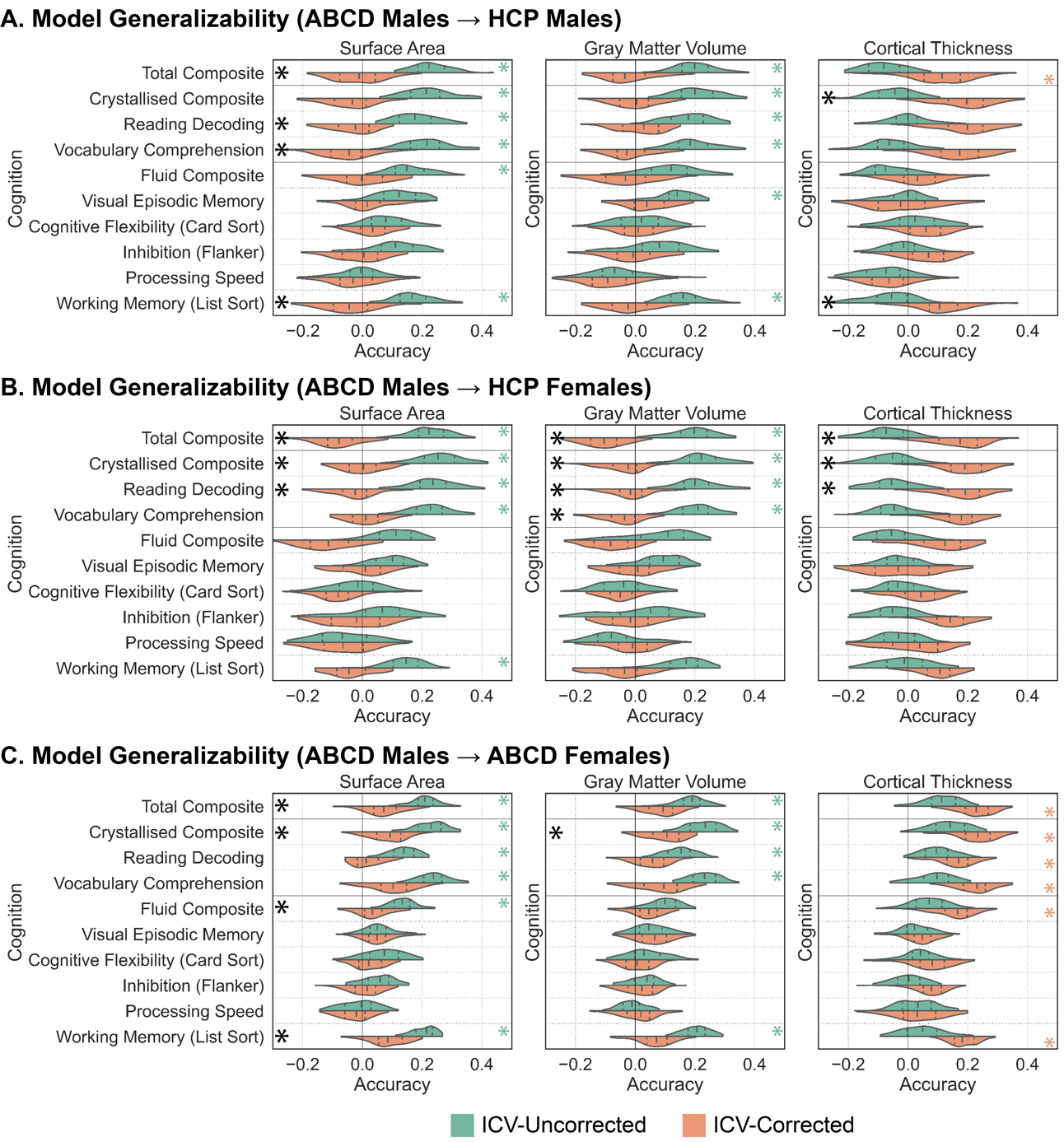

Figure S8: **Accounting for intracranial volume differentially impacts model generalizability in terms of predictive accuracies across surface area, gray matter volume, and cortical thickness in a sex specific manner.** Prediction accuracies (Pearson’s correlation coefficient between observed and predicted scores) for sex-specific models trained on ABCD males to predict cognitive scores and evaluated on HCP males (A), HCP females (B), and ABCD females (C). Predictions based on surface area (left), gray matter volume (middle), and cortical thickness (right) using raw (green) and ICV proportion-corrected (orange) anatomical properties are shown. Green and orange asterisks (*) denote that the model performed above chance levels based on permutation tests (corrected p<0.05). Black asterisks (*) denote that model performance was significantly different between the raw and ICV proportion-corrected predictions based on exact tests for differences (corrected p<0.05). The shape of the violin plots indicates the entire distribution of values, dashed lines indicate the median, and dotted lines indicate the interquartile range.

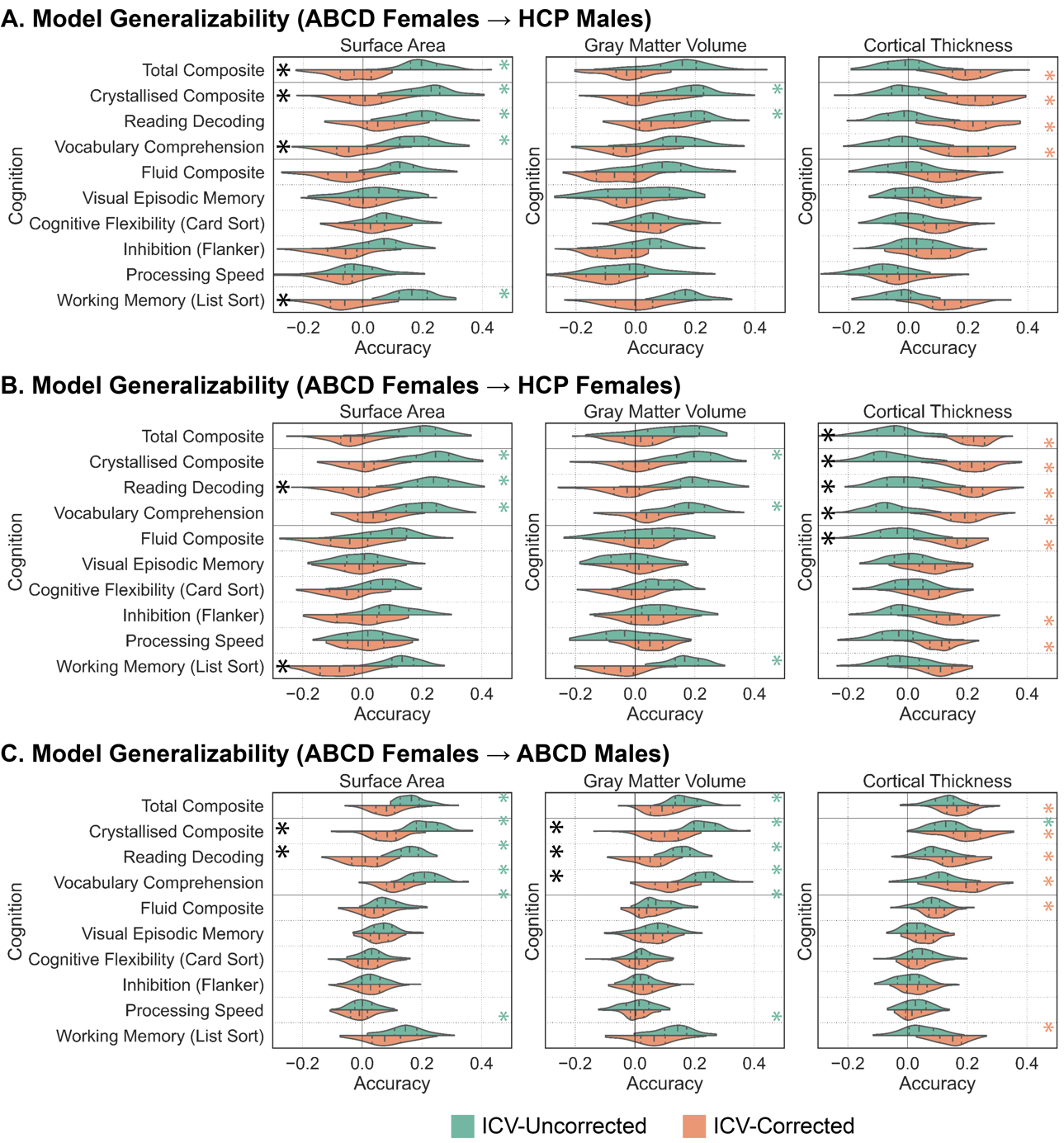

Figure S9: **Accounting for intracranial volume differentially impacts model generalizability in terms of predictive accuracies across surface area, gray matter volume, and cortical thickness in a sex specific manner.** Prediction accuracies (Pearson’s correlation coefficient between observed and predicted scores) for sex-specific models trained on ABCD females to predict cognitive scores and evaluated on HCP males (A), HCP females (B), and ABCD males (C). Predictions based on surface area (left), gray matter volume (middle), and cortical thickness (right) using raw (green) and ICV proportion-corrected (orange) anatomical properties are shown. Green and orange asterisks (*) denote that the model performed above chance levels based on permutation tests (corrected p<0.05). Black asterisks (*) denote that model performance was significantly different between the raw and ICV proportion-corrected predictions based on exact tests for differences (corrected p<0.05). The shape of the violin plots indicates the entire distribution of values, dashed lines indicate the median, and dotted lines indicate the interquartile range.

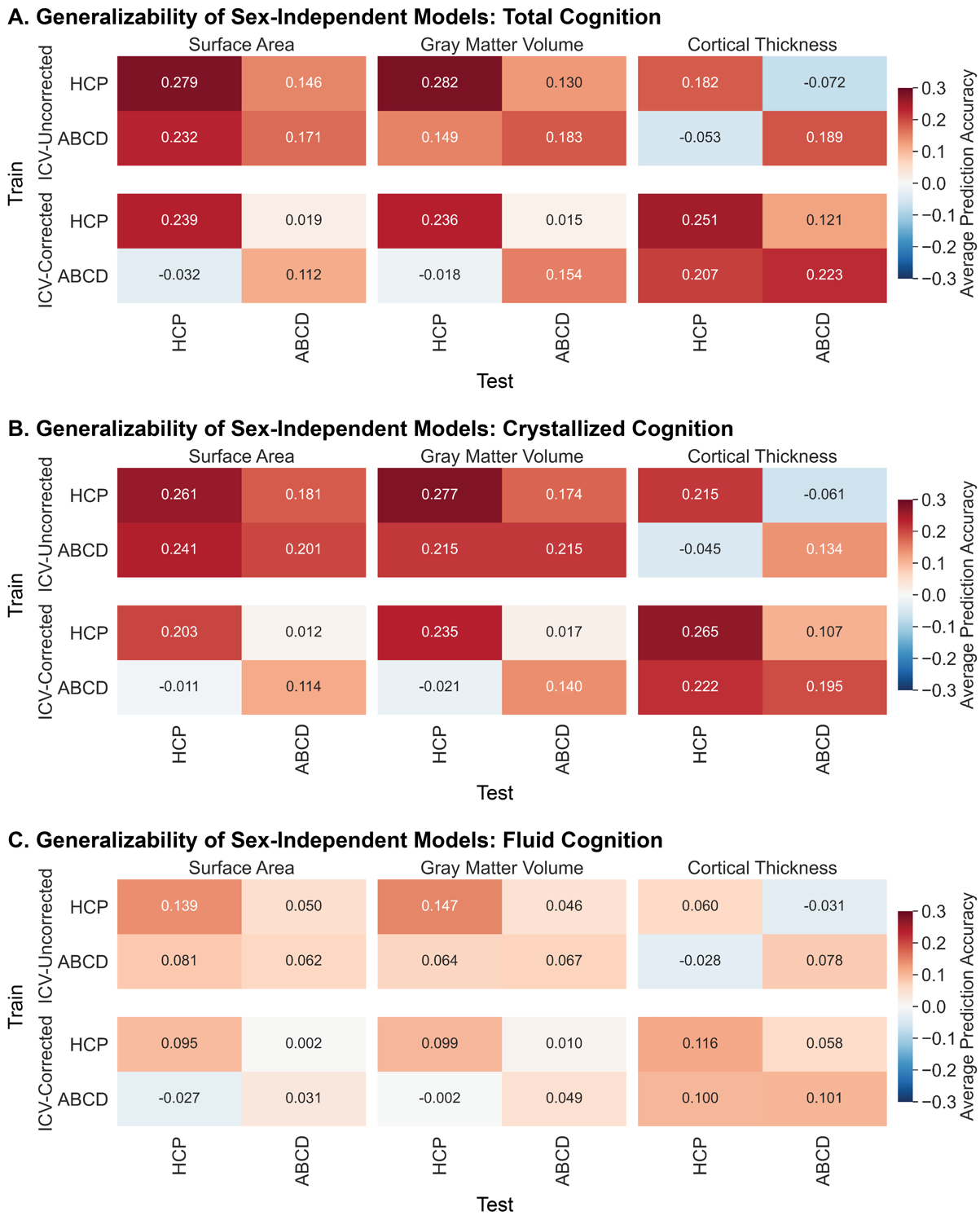

Figure S10: **Intracranial volume correction reduces generalizability of sex-independent models based on surface area and gray matter volume but increases generalizability of models based on cortical thickness.** Generalizability of sex-independent models across sexes (males and females) and datasets (HCP and ABCD) to predict the Total Composite score (A), Crystallized Composite and associated task scores (B), and Fluid Composite and associated task scores (V). Average prediction accuracies based on surface area (left), gray matter volume (middle), and cortical thickness (right) using raw anatomical properties are shown in the top panels, and predictions using ICV proportion-corrected anatomical properties are shown in the bottom panels. The populations that the models were trained on are shown along the rows, and the populations that the models were tested on are shown along the columns.

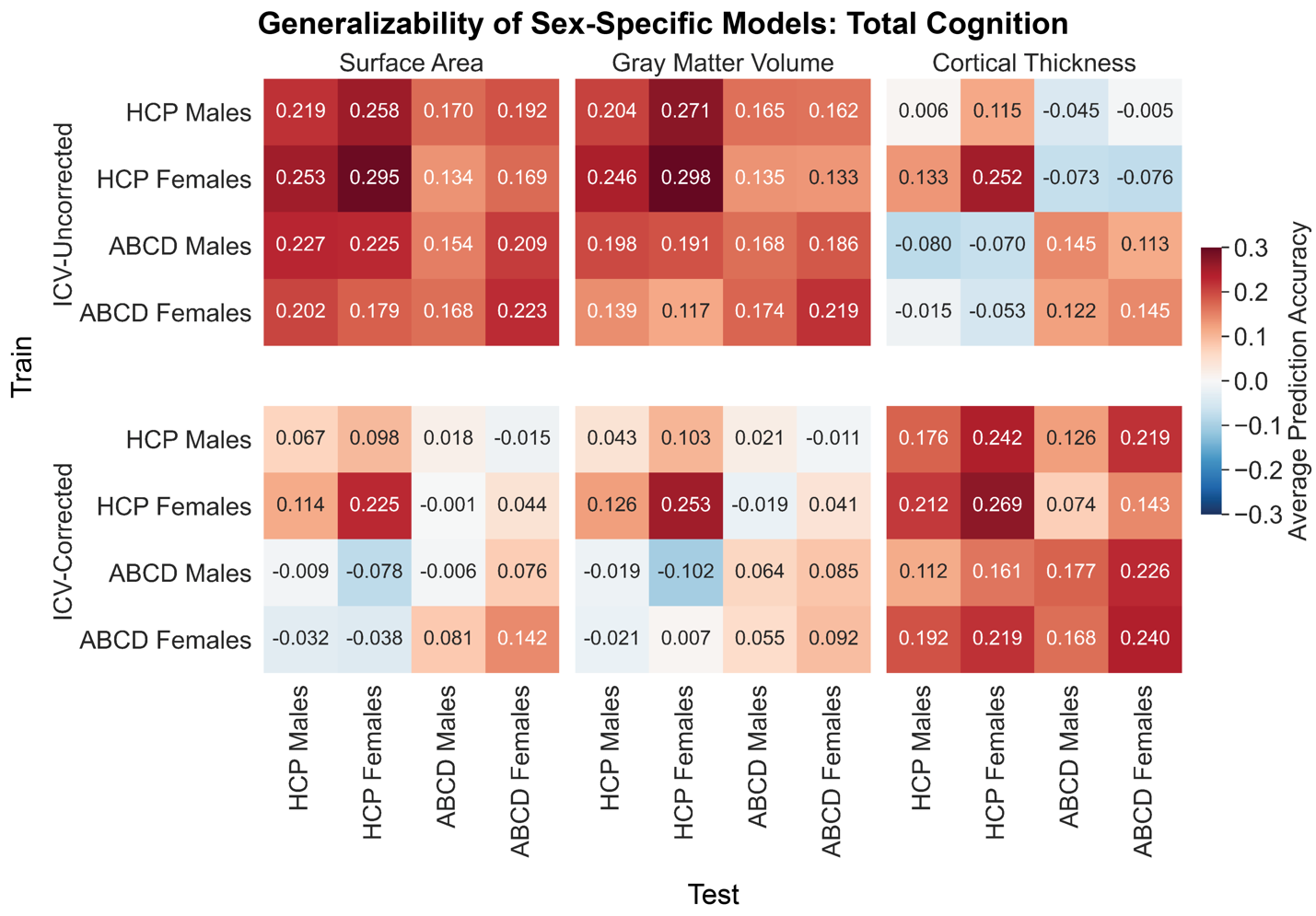

Figure S11: **Intracranial volume correction reduces generalizability of sex-specific models based on surface area and gray matter volume but increases generalizability of models based on cortical thickness to predict total Cognition.** Generalizability of sex-independent models across sexes (males and females) and datasets (HCP and ABCD). Prediction accuracies to predict the Total Cognition Composite score based on surface area (left), gray matter volume (middle), and cortical thickness (right) using raw anatomical properties are shown in the top panels, and predictions using ICV proportion-corrected anatomical properties are shown in the bottom panels. The populations that the models were trained on are shown along the rows, and the populations that the models were tested on are shown along the columns.

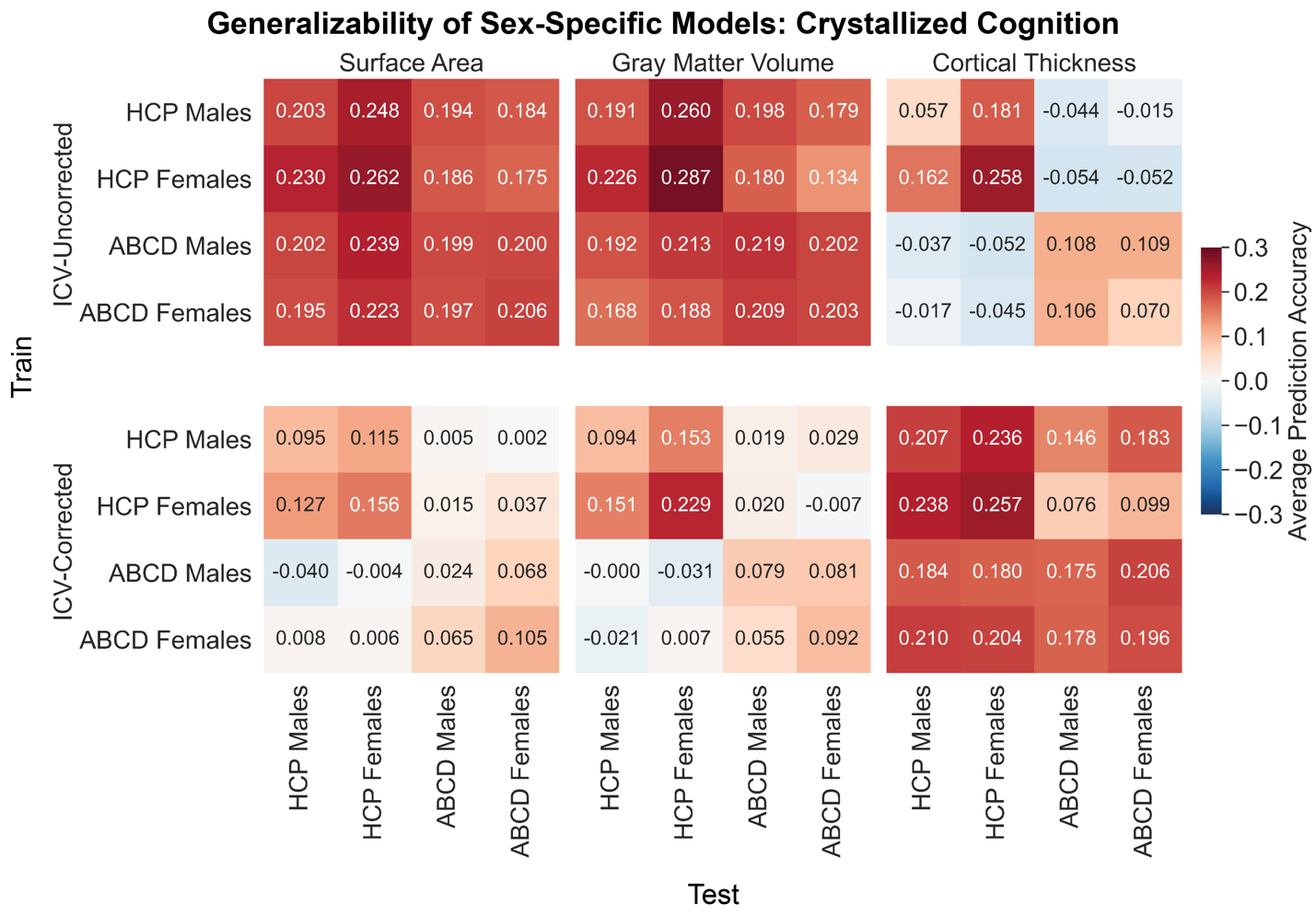

Figure S12: **Intracranial volume correction reduces generalizability of sex-specific models based on surface area and gray matter volume but increases generalizability of models based on cortical thickness to predict crystallized abilities.** Generalizability of sex-independent models across sexes (males and females) and datasets (HCP and ABCD). Average prediction accuracies across the crystallized composite score and individual task scores within the crystallized domain based on surface area (left), gray matter volume (middle), and cortical thickness (right) using raw anatomical properties are shown in the top panels, and predictions using ICV proportion-corrected anatomical properties are shown in the bottom panels. The populations that the models were trained on are shown along the rows, and the populations that the models were tested on are shown along the columns.

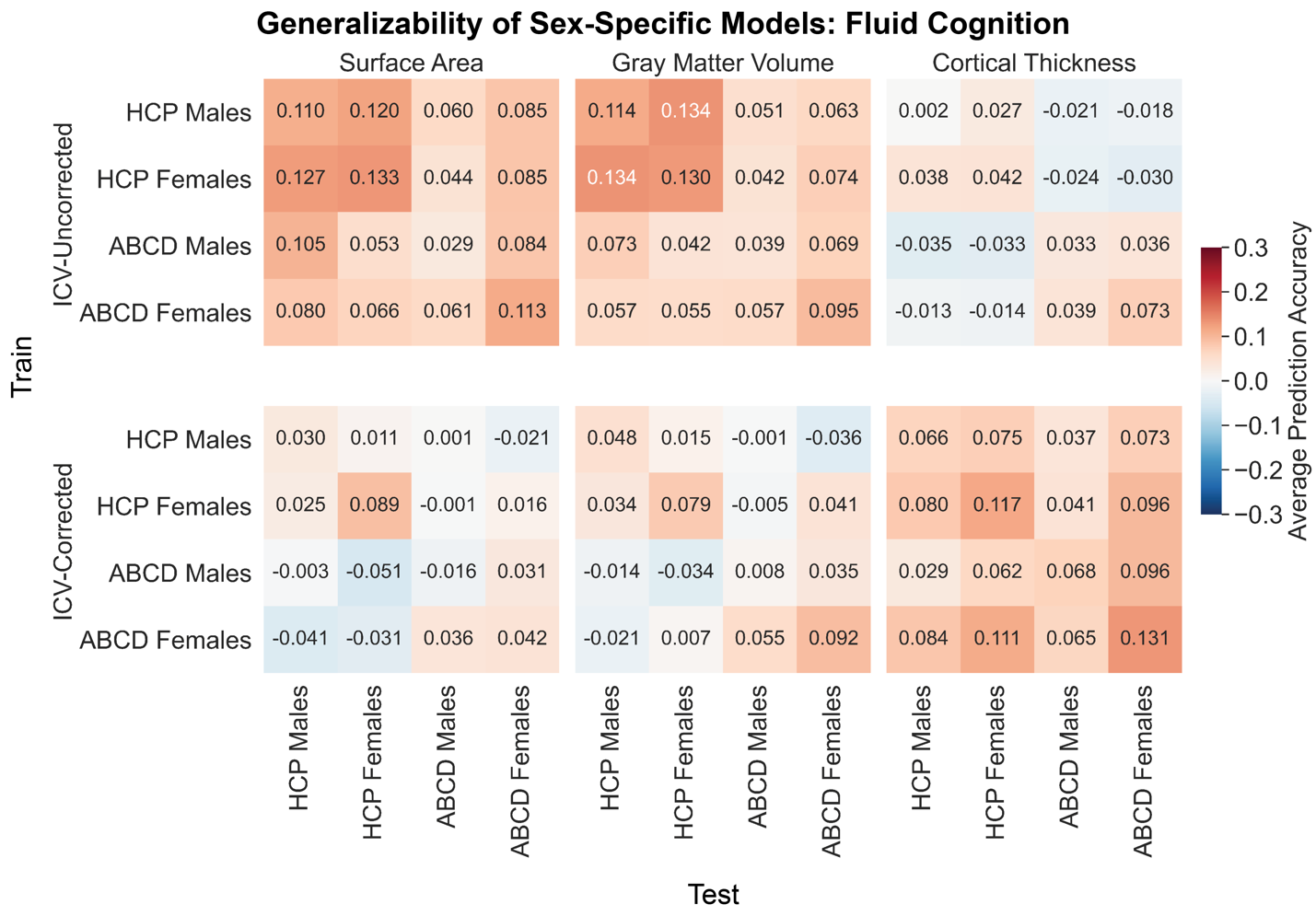

Figure S13: **Intracranial volume correction reduces generalizability of sex-specific models based on surface area and gray matter volume but increases generalizability of models based on cortical thickness to predict fluid abilities.** Generalizability of sex-independent models across sexes (males and females) and datasets (HCP and ABCD). Average prediction accuracies across the fluid composite score and individual task scores within the fluid domain based on surface area (left), gray matter volume (middle), and cortical thickness (right) using raw anatomical properties are shown in the top panels, and predictions using ICV proportion-corrected anatomical properties are shown in the bottom panels. The populations that the models were trained on are shown along the rows, and the populations that the models were tested on are shown along the columns.

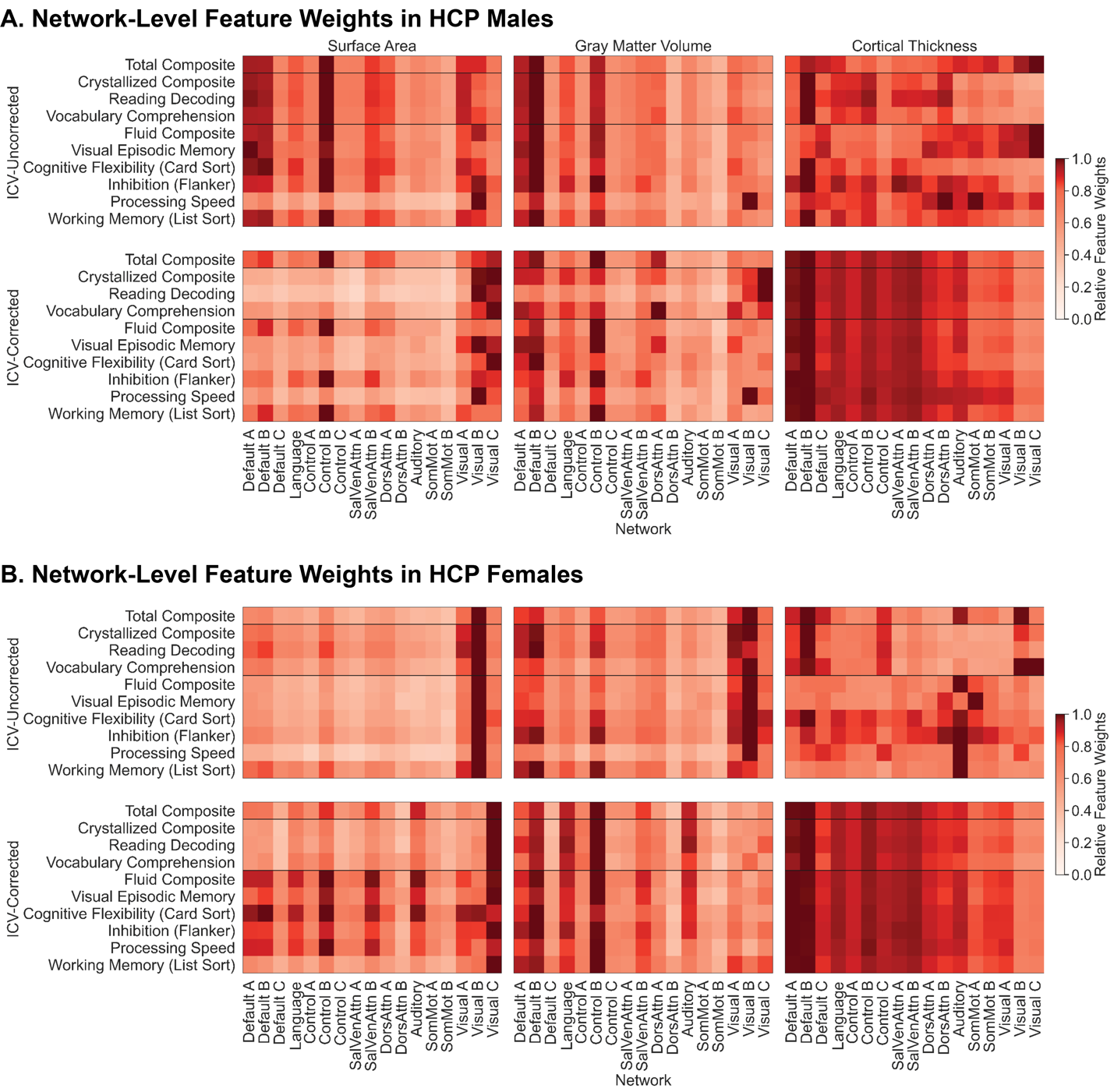

Figure S14: **The predictive relationships linking cognition with the anatomy of association and unimodal cortices in adults can be revealed or obscured though the use of intracranial volume correction.** Absolute relative network-level sex-specific feature weights to predict each of the cognitive scores in HCP males (A) and HCP females (B). Feature weights for models based on surface area (left), gray matter volume (middle), and cortical thickness (right) using raw anatomical properties are shown in the top panels, and predictions using ICV proportion-corrected anatomical properties are shown in the bottom panels. SalVenAttn – Salience/Ventral Attention; DorsAttn – Dorsal Attention; SomMot – Somatomotor. Networks are ordered from heteromodal (left) to unimodal (right).

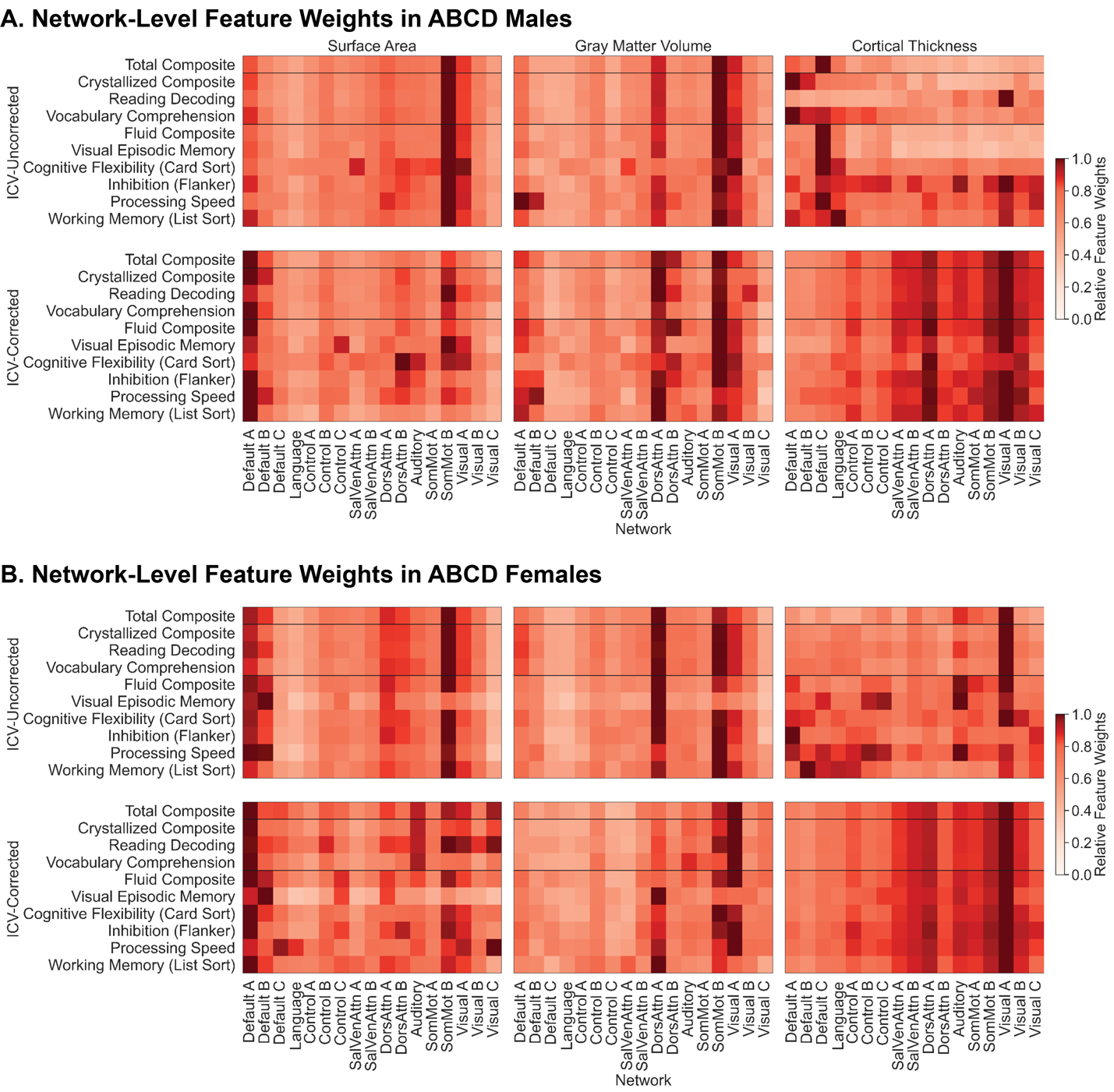

Figure S15: **The predictive relationships linking cognition with the anatomy of association and unimodal cortices in children can be revealed or obscured though the use of intracranial volume correction.** Absolute relative network-level sex-specific feature weights to predict each of the cognitive scores in ABCD males (A) and ABCD females (B). Feature weights for models based on surface area (left), gray matter volume (middle), and cortical thickness (right) using raw anatomical properties are shown in the top panels, and predictions using ICV proportion-corrected anatomical properties are shown in the bottom panels. SalVenAttn – Salience/Ventral Attention; DorsAttn – Dorsal Attention; SomMot – Somatomotor. Networks are ordered from heteromodal (left) to unimodal (right).
